# Obligate intracellular *Orientia tsutsugamushi* impedes *TP53* expression to inhibit DNA damage-induced apoptosis

**DOI:** 10.1101/2025.04.07.647604

**Authors:** Paige E. Allen, David L. Armistead, Svetlana Blinova, Jason A. Carlyon

## Abstract

Infections by intracellular pathogens often cause insult to host cell DNA, which stimulates responses that ultimately eliminate the damaged cell and hence the microbial niche. p53 is an innate immunity mediator that responds to DNA damage and intracellular infection by transcriptionally activating pathways that arrest the cell cycle, repair DNA, and elicit apoptosis. How pathogens counter p53 are incompletely understood. Here, we demonstrate that the endotheliotropic obligate intracellular bacterium *Orientia tsutsugamushi* blocks expression of *TP53* by targeting its upstream region to nearly deplete p53 levels. Contrary to the unrestricted proliferation expected based on the transcriptome of p53-deficient infected cells, *Orientia* arrests the cell cycle at S phase to promote bacterial replication. It protects host endothelial cells from DNA damage even if induced by etoposide and delays genotoxic-dependent apoptosis until late in infection once a high bacterial load has been achieved. *TP53* downregulation, protection against genotoxicity, and inhibition of DNA damage-dependent apoptosis are executed by the *Orientia* nucleomodulatory effector, Ank13. Therefore, *O. tsutsugamushi* inhibits *TP53* expression and genotoxicity to reconfigure the intracellular environment of its host cell into one that favors bacterial replication.

## INTRODUCTION

Intracellular pathogens reshape the host cellular environment into one that supports optimal replication. Intracellular infections are often genotoxic, which elicits DNA damage responses that ultimately trigger apoptosis to eliminate the marred cell along with its microbial inhabitants^1,2^. As a countermeasure, intracellular microbes must prevent host genome instability and inhibit programmed cell death responses to sustain their niches^2^. The pathogenic mechanisms that facilitate this evolutionarily conserved survival strategy are inadequately defined.

Known as “the guardian of the genome”, p53 is a tumor suppressing transcription factor that is conserved among higher eukaryotes and controls the response to DNA damage^2,3^. Its importance to maintaining cellular homeostasis is underscored by the fact that, *TP53*, the gene that encodes p53, is mutated in more than half of human cancers. Under normal conditions, p53 is marked for degradation by the HDM2 (human double minute 2) E3 ubiquitin ligase. When DNA damage is incurred, p53 is stabilized and activates a transcriptional program that temporarily arrests the cell cycle to allow for DNA repair. If the damage is irreversible, p53 induces senescence, a state of permanent cell cycle arrest where the cell remains viable, or apoptosis^1,2^. It is also a central mediator of innate immunity to intracellular microbes^4^. p53 is therefore an antimicrobial hurdle that these organisms must overcome. Several intracellular pathogens orchestrate p53 degradation^5–19^. Multiple viruses inhibit p53 translation while the cattle parasite, *Theileria annulata*, blocks p53 translocation into the nucleus^20–26^. *TP53* is post-transcriptionally repressed by a TLR4-NF-κB-mediated cellular pathway activated in response to lipopolysaccharides from *Klebsiella pneumoniae* and other Enterobacteria^27^. Microbial p53 suppression has been repeatedly linked to tumorigenesis^1,5–30^. No pathogen has been shown to directly inhibit *TP53* expression.

*Orientia tsutsugamushi* is a mite-transmitted obligate intracellular bacterium that causes scrub typhus, an emerging spotted-fever illness and the deadliest human rickettsiosis ^31,32^. Scrub typhus is endemic throughout the Asia-Pacific and occurs in numerous African, Middle Eastern, and South American countries^32–37^. *Orientia* recently was detected in mites in North Carolina potentially extending the disease’s geographic footprint to North America^38,39^. Diagnosis can be difficult, treatment options are limited, and there is no vaccine^40,41^. Scrub typhus initiates as an acute non-specific febrile illness accompanied by localized vasculitis and has not been linked to cancer. If left untreated or if antibiotic therapy is delayed, severe symptoms including respiratory distress, renal failure, meningoencephalitis, and myocarditis ensue. Moreover, widespread vasculitis can lead to hypotensive shock, multiorgan failure, and death^41,42^. Scrub typhus vasculitis is linked to heavy *O. tsutsugamushi* burdens of endothelial cells^43–48^. Following endothelial cell invasion, the pathogen replicates to high numbers in the cytosol^49–53^. By around 72 h post-infection (p.i.) logarithmic replication has ceased and the cell is loaded with bacteria primed for release^50,51,53,54^. At this timepoint, DNA damage and apoptosis are first observable, both in infected endothelial cell lines and the endothelium of infected mice^43,52,55^. High *O. tsutsugamushi* loads in endothelial cells are also associated with an overzealous inflammatory response that further contributes to the vasculitis and other serious manifestations^43,45–48^. These observations suggest that *O. tsutsugamushi* inhibits DNA damage-induced apoptosis to replicate sufficiently.

Here, we report that *O. tsutsugamushi* inhibits *TP53* expression by targeting its upstream region to severely reduce p53 levels. Infected endothelial cells exhibit transcriptional profiles consistent with a lack of p53. However, contrary to the uncontrolled proliferation anticipated for such cells, *O. tsutsugamushi* pauses the cell cycle at S phase to maximize its intracellular growth. It protects endothelial cells from genotoxicity and inhibits DNA damage-induced apoptosis until 72 h p.i. The bacterium’s abilities to downregulate *TP53* by targeting its upstream region, inhibit DNA damage, and delay genotoxic-dependent apoptosis involves its nucleomodulin, Ank13. These modulatory functions depend on the effector’s nucleotropism and ability to coopt the host SCF E3 ubiquitin ligase complex. Our findings outline a new mechanism by which a pathogen modulates p53 and its downstream responses to increase host cell longevity and optimize intracellular infection.

## RESULTS

### O. tsutsugamushi downregulates TP53

The *O. tsutsugamushi* Ikeda strain, which was originally recovered from a scrub typhus patient in Japan, causes severe disease in humans and is fatal in laboratory mice^56–60^. Ikeda was used for all experiments and will hereafter be referred to as *O. tsutsugamushi* or *Orientia* unless discussed in the context of other strains. The endothelial cell pathways that *O. tsutsugamushi* infection alters at the transcriptional level are poorly characterized. RNA-seq was performed on paired infected and uninfected EA.hy926 human vascular endothelial cells at 8, 48, and 72 h p.i., timepoints that correspond to lag, mid-exponential, and late-exponential *Orientia* replication, respectively (Supplementary Fig. 1a)^50,53,61^. Three biological replicates with similar bacterial loads were evaluated (Supplementary Fig. 1b). Pearson’s correlation coefficient and principal component analyses confirmed that differentially expressed genes (DEGs) could be defined at each timepoint (Supplementary Fig. 1c, d). DEGs were defined as having a -log_10_(padj) value of ≥ 1.3 and a log_2_(fold change) of ≥ 1 or ≤ -1 in infected compared to uninfected cells (Supplementary Data 1).

At 8, 48, and 72 h p.i., DEGs made up approximately 1%, 13%, and 30% of transcripts detected, respectively (Fig. 1a-c). DEGs were predominantly upregulated at 8 and 48 h p.i., but comparably up- and downregulated at 72 h p.i. (Fig. 1a-c). Only four DEGs were altered at all timepoints (Fig. 1a-d). *TP53,* which encodes p53^3^, and *CLIC1,* which contributes to intracellular physiological stability^62^, were downregulated. *BST2* and *MX2,* which promote antiviral responses^63^ and innate immunity^64^, respectively, were upregulated.

**Fig. 1.**
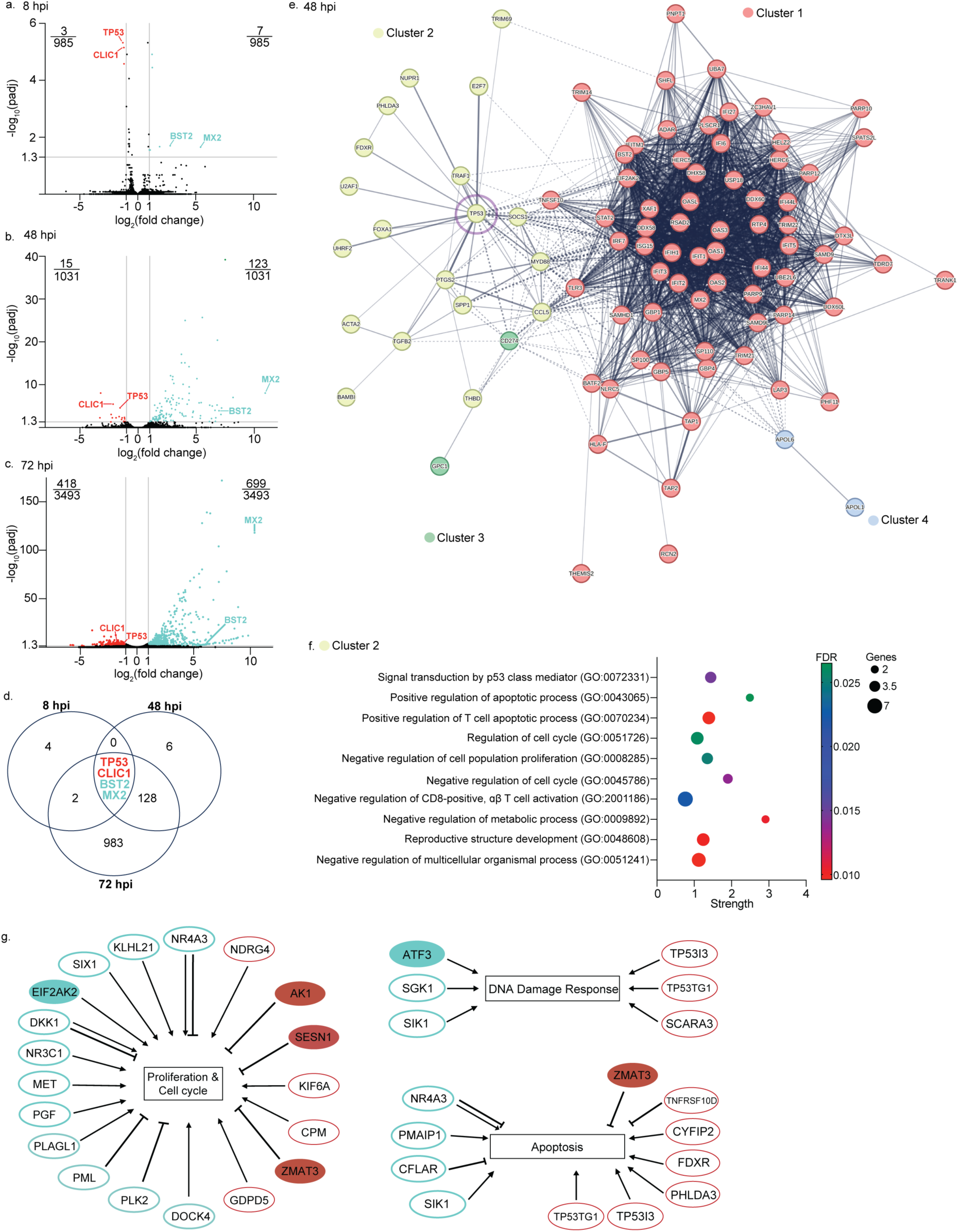
*O. tsutsugamushi* transcriptionally modulates *TP53* and downstream pathways. **(a-c)** Volcano plots of the gene expression profiles for *O. tsutsugamushi* infected EA.hy926 cells compared to uninfected cells. Each dot corresponds to an individual gene. Teal and red dots indicate differentially expressed genes (DEGs) that are up- or downregulated, respectively. Black dots represent genes that are not differentially expressed. Grey lines delineate DEGs as genes with a significant -log_10_(padj)value (horizontal lines > 1.3) and log_2_(fold change) (vertical lines > 1 or < -1). Fractions indicate the numbers of DEGs up- (upper right) or downregulated (upper left) out of the total number of genes identified. **(d)** Venn diagram showing the number of unique and shared DEGs at 8 h p.i. (10 total), 48 h p.i. (138 total), and 72 h p.i. (1117 total). Overlapping regions display the number of common DEGs among intersecting groups with *TP53*, *CLIC1*, *BST2*, and *MX2* being present at all timepoints and colored to indicate up- (teal) or downregulated (red) expression. **(e)** STRING analysis of all DEGs at 48 h p.i. Disconnected nodes were removed, and Markov Clustering was employed to define prominent clusters. Solid lines denote a direct connection between genes. Line thickness indicates the strength of the connection based on existing literature. Dotted lines represent connections between genes that are on the cusp of multiple clusters. Four clusters are defined: Cluster 1 (red) – interferon signaling and ISG15-protein conjugation; Cluster 2 (yellow) – p53-related pathways (*TP53* indicated by a purple circle); Cluster 3 (green) – signaling receptors; Cluster 4 (blue) – apolipoproteins. **(f)** Bubble plot showing the top 10 GO terms subdivided by biological processes that are associated with Cluster 2. Bubble size correlates with the number of DEGs from the input present in that GO term, bubble color indicates the false discovery rate (FDR), and bubble position on the x-axis shows the strength of the enrichment. **(g)** Schematic of up- (teal) or downregulated (red) p53-associated DEGs that negatively (inhibitor lines) or positively (arrows) modulate proliferation and cell cycle, DNA damage response, and apoptosis. Filled in ovals indicate proteins that directly bind p53.

DEGs (Supplementary Data 1) were subjected to STRING (Search Tool for the Retrieval of Interacting Genes/Proteins) analysis with Markov clustering to identify host pathways altered at the transcriptional level during infection (Fig. 1e; Supplementary Fig. 2). At 48 h p.i., four distinct clusters were observed. Cluster 1 contained genes associated with interferon signaling and ISG15-(interferon stimulated gene 15) protein conjugation and produced gene ontology (GO) terms centered around antimicrobial responses (Fig. 1e; Supplementary Data 2). Cluster 2 expanded from *TP53* with the top 10 GO terms related to p53-regulated pathways (Fig. 1e, f; Supplementary Data 2). Clusters 3 and 4 contained only two DEGs that encoded signaling receptors and apolipoproteins, respectively. *TP53* also shared a connection with DEGs in Clusters 1 and 3 (Fig. 1e; Supplementary Data 2). At 72 h p.i., DEGs were enriched in pathways involved in p53-related DNA damage responses; interferon, IL-10, and NF-κB signaling; eukaryotic translation elongation; the mitotic spindle checkpoint; and the inflammasome (Supplementary Fig. 2; Supplementary Data 2). Cross-referencing DEGs from all timepoints to a comprehensive list of known p53 target genes^65^ identified a total of 76 that exhibited expression patterns indicative of an absence of p53 (Supplementary Data 3; Supplementary Fig. 3)^2,3,66–69^. Of these, 31 aligned with unrestricted proliferation and cell cycle progression, limited DNA damage repair responses, and inhibition of apoptosis (Fig. 1g; Supplementary Fig. 3). Thus, *O. tsutsugamushi* downregulates *TP53* expression, which alters multiple p53-regulated pathways at the transcriptional level.

### *O. tsutsugamushi* rapidly reduces p53 levels

RT-qPCR analysis validated that *TP53* expression is downregulated throughout *Orientia* infection (Fig. 2a). Immunoblot studies detected a sustained near absence of p53 as early as 4 h p.i. (Fig. 2b-e). The *O. tsutsugamushi* immunodominant outer membrane protein, TSA56 (56-kDa type-specific antigen), served as an infection control while human glyceraldehyde-3-phosphate dehydrogenase (GAPDH) was a loading control for all western blots. Chloramphenicol, an antibiotic that crosses eukaryotic membranes to inhibit bacterial protein synthesis and hence replication, prevented *O. tsutsugamushi* from lowering p53 levels (Fig. 2f, g). Expression of p21, a p53 target that regulates cell cycle arrest, senescence, and apoptosis^70^, was reduced throughout infection as early as 24 h p.i. and in a chloramphenicol-sensitive manner (Supplementary Fig. 4). Additionally, scrub typhus clinical isolates UT76, Karp, and Gilliam grow in EA.hy926 cells comparably to Ikeda and impede *TP53* and p53 expression throughout infection (Supplementary Fig. 5). Therefore, the conserved ability among *O. tsutsugamushi* strains to rapidly downregulate *TP53* expression correlates with a sustained reduction in p53 that has downstream consequences.

**Fig. 2.**
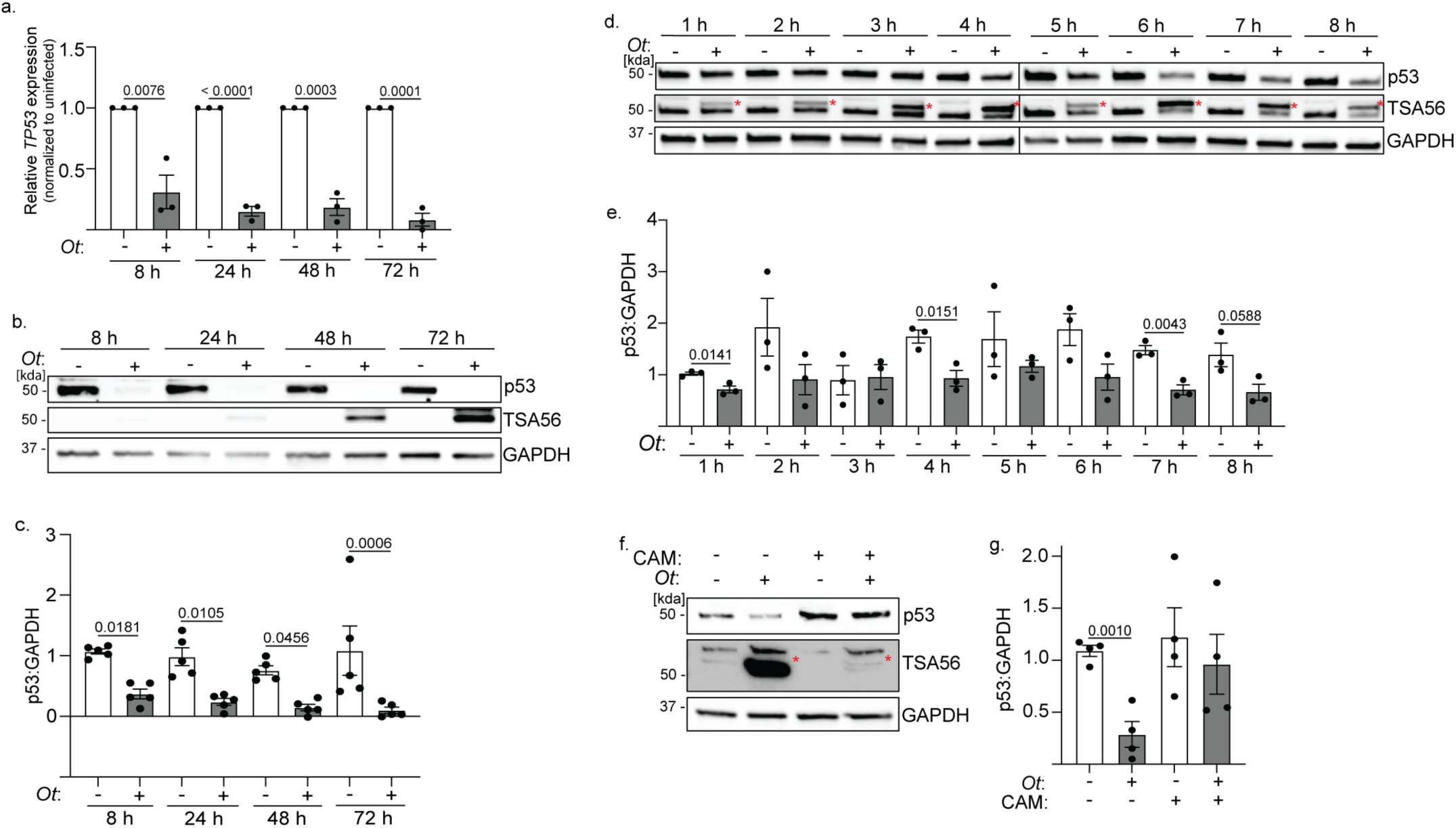
*O. tsutsugamushi* significantly reduces p53 levels. **(a)** RNA and **(b-g)** whole cell lysates (WCLs) were collected from uninfected and *O. tsutsugamushi* infected EA.hy926 cells at **(a-c)** 8, 24, 48, and 72 h p.i. or **(d, e)** every hour for 8 h p.i. and subjected to **(a)** RT-qPCR or **(b**, **d)** western blot and **(c, e)** densitometry analysis. **(f**, **g)** Uninfected and infected EA.hy926 cells were treated with 34 µg/mL of chloramphenicol (CAM) or vehicle (VEH) control at 2 h p.i. At 72 h p.i., WCLs were collected and **(f)** western blot and **(g)** densitometry analysis was performed. Blots were probed with antibodies against p53, TSA56, or GAPDH. Red asterisks (*) denote TSA56 in infected samples. Mean normalized ratios ± SEM of **(a)** relative *TP53* gene expression normalized to *GAPDH* or **(c, e, g)** p53:GAPDH were calculated (*n* = 3-5 independent experiments). Statistical significance was evaluated by One-way ANOVA followed by Tukey’s *post hoc* test. Statistically significant values (P ≤ 0.05) are displayed.

### *O. tsutsugamushi* specifically targets *TP53* gene expression

p53 is constitutively expressed to ensure a rapid response to stress. This is counterbalanced by the ubiquitin ligase, HDM2, which polyubiquitinates p53 to target it for proteasomal degradation^16,27^. Many intracellular pathogens exploit HDM2 to degrade p53^5–19^. We investigated if *Orientia* employs a similar mechanism to target p53. HDM2 transcript and protein were pronouncedly lowered throughout infection (Fig. 3a-c; Supplementary Data 1), suggesting that HDM2 is inessential for *O. tsutsugamushi* to modulate p53 levels. To confirm that HDM2 activity is unnecessary for *Orientia* fitness, Nutlin 3a, a steric inhibitor that prevents HDM2 ubiquitination of p53^71^, was employed to stabilize p53 (Supplementary Fig. 6a). Nutlin 3a had no effect on the *O. tsutsugamushi* yield (Fig. 3d). Next, p53 levels were assessed in infected and uninfected cells treated with the proteasome inhibitor, bortezomib. Proteasome inhibition was confirmed by increased Nrf1 expression (Fig. 3e)^72^. p53 levels were lower in infected cells regardless of bortezomib treatment (Fig. 3e, f). Thus, *O. tsutsugamushi* inhibits p53 expression independent of HDM2 and the proteasome.

**Fig. 3.**
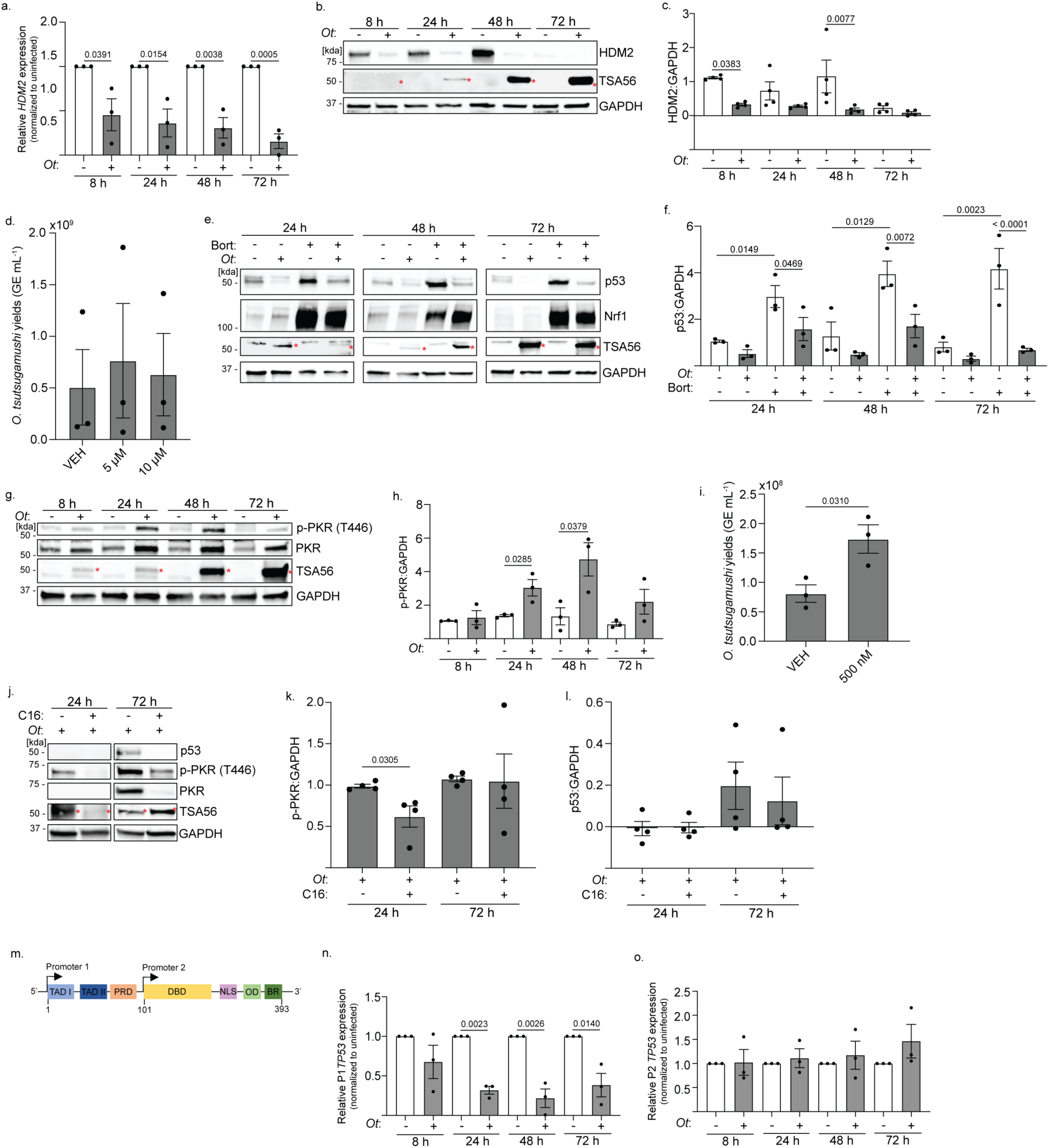
*O. tsutsugamushi* specifically targets *TP53* gene expression. **(a-c)** RNA and WCLs were collected from uninfected and *O. tsutsugamushi* infected EA.hy926 cells at 8, 24, 48, and 72 h p.i. and subjected to **(a)** RT-qPCR, or **(b)** western blot and **(c)** densitometry analysis. Blots were probed with antibodies against HDM2, TSA56, or GAPDH. **(a)** Relative *HDM2* gene expression normalized to *GAPDH* or **(c)** HDM2:GAPDH were calculated. **(d)** The *O. tsutsugamushi* yield was determined for vehicle (VEH) or Nutlin 3a (Nut3a) treated infected EA.hy926 cells. **(a, c, d)** Data are means ± SEM (*n* = 3-4 independent experiments). **(e-f)** Uninfected or infected EA.hy926 cells were treated with VEH or bortezomib (Bort; 50 nM) for 24 h prior to collection and **(e)** western blotting. Blots were probed for p53, Nrf1, TSA56, or GAPDH. Mean normalized ratios ± SEM of **(f)** p53:GAPDH were determined by densitometry (*n* = 3 independent experiments). **(g, h)** WCLs were collected from uninfected and *O. tsutsugamushi* infected EA.hy926 cells and subjected to **(g)** western blot. Blots were probed with antibodies against phosphorylated (p)-PKR (T446), PKR, TSA56, or GAPDH. Mean normalized ratios ± SEM of **(h)** p-PKR:GAPDH were determined by densitometry (*n* = 3 independent experiments). **(i-l)** *O. tsutsugamushi* infected EA.hy926 cells were treated with VEH or C16 (100 nM). **(i)** Lysates were collected at 72 h p.i. for qPCR to determine *O. tsutsugamushi* GE mL^-1^. **(j-l)** WCLs were collected at 24 and 72 h p.i. for **(j)** western blot and **(k, l)** densitometry analysis. Blots were probed with antibodies against p53, p-PKR (T446), PKR, TSA56, or GAPDH. Mean normalized ratios ± SEM of **(k)** p-PKR:GAPDH and **(l)** p53:GAPDH were calculated (*n* = 3 independent experiments). **(m)** Schematic of *TP53* and its promoters. **(n, o)** RNA was collected from uninfected and *O. tsutsugamushi* infected EA.hy926 cells and subjected to RT-qPCR. Mean normalized ratios ± SEM of relative *TP53* **(n)** P1 and **(o)** P2 gene expression normalized to *GAPDH* (*n* = 3 independent experiments). Red asterisks (*) denote TSA56 in infected samples. Statistical significance was evaluated by One-way ANOVA followed by Tukey’s *post hoc* test. Statistically significant values (P < 0.05) are displayed.

Double-stranded RNA viruses trigger the protein kinase R (PKR) stress response that inhibits host cell translation and thereby p53 expression^73,74^. PKR is activated by phosphorylation of T446^75^. During *O. tsutsugamushi* infection, PKR activation was significantly increased at 24 and 48 h p.i. (Fig. 3g, h). Inhibiting PKR phosphorylation using the small molecule inhibitor, C16^76^, failed to restore p53 levels in infected cells and even led to a higher bacterial load (Fig. 3i-l). Therefore, despite activating the PKR stress response, *Orientia* blocks p53 expression in a PKR-independent manner.

*TP53* encodes two families of p53 isoforms via alternative promoters, P1 and P2 (Fig. 3m). Canonical *TP53* transcription initiated from P1 generates an mRNA that is translated into full-length p53. Non-canonical transcription from P2 produces a transcript encoding either of two truncated isoforms^77^. P1 and P2 encoded *TP53* mRNAs share a 3’-UTR that is critical for their stability. Lipopolysaccharides of enterobacteria with tumorigenic potential destabilize both P1 and P2 *TP53* mRNA at the 3’-UTR, which decreases p53 levels and p53-regulated responses^27^. RT-qPCR conducted with isoform-specific primers found that *O. tsutsugamushi* prevents *TP53* expression from P1 but not P2 (Fig. 3n, o), suggesting that it employs the novel mechanism of targeting the P1 *TP53* isoform at its upstream region to block full-length p53 production.

### *Orientia* modulates the cell cycle

p53 deficiency typically leads to uncontrolled proliferation^66–68^, a phenotype that is suggested by the transcriptome of *O. tsutsugamushi* infected endothelial cells (Supplementary Data 3; Figure 1g; Supplementary Figure 3). Accordingly, we assessed proliferation rates of uninfected and infected EA.hy926 cells that had been synchronized at G0/G1 by serum starvation and released when *Orientia* was added. Immunofluorescence microscopy was used to confirm that ≥ 75% of the cells in infected cultures harbored bacteria (Fig. 4a, b). Starting at 24 h, proliferation rates of infected cultures trended lower and were significantly reduced by two-fold at 48 h (Fig. 4c, d). Hence, despite p53 inhibition, infected cell proliferation is hindered and is most pronouncedly so during *O. tsutsugamushi* mid-log growth. We then measured cyclin protein turnover to monitor cell cycle progression (Fig. 4e)^68^. Cyclin D1 and cyclin D3 levels were robustly reduced during infection in a chloramphenicol-sensitive manner (Fig. 4f-h; Supplementary Fig. 7a-c), which indicates cell cycle progression through G1 and correlates with the bacterial protein synthesis-dependent decrease in p53 (Fig. 2f, g)^68^. In contrast, cyclin E levels were significantly higher at 72 h p.i. (Fig. 4f, i). Cyclin A was slightly elevated at 24 and 48 h p.i. (Fig. 4f, j), while cyclin B1 levels trended lower in infected cells at all time points (Fig. 4f, k). These data indicate that *O. tsutsugamushi* promotes a pause in the host cell cycle at S phase during its exponential replication.

**Fig. 4.**
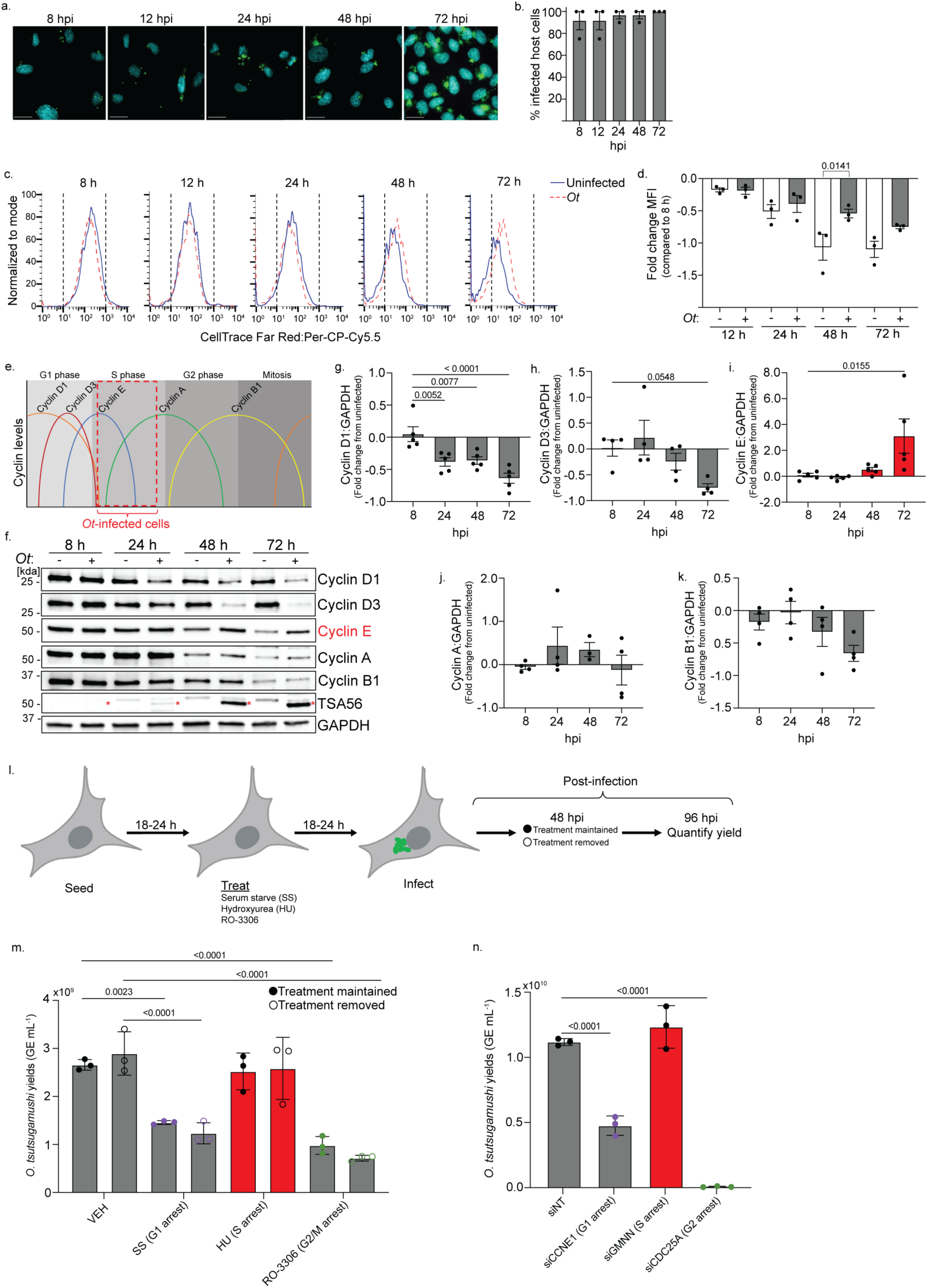
*O. tsutsugamushi* modulates the cell cycle as a promicrobial strategy. **(a-d)** CellTrace host cell proliferation assay. **(a)** Coverslips with *O. tsutsugamushi* infected EA.hy926 cells were methanol fixed and immunolabeled for TSA56 (green), stained with DAPI (blue), and imaged using immunofluorescent microscopy. **(b)** The percentage of infected cells was determined by examining 100 cells at each timepoint. **(c)** Host cell proliferation was assessed using flow cytometry. **(d)** The fold change in mean fluorescent intensity (MFI) was determined by normalizing each sample to the 8 h timepoint. **(e)** Schematic of cyclin protein dynamics at different phases of the host cell cycle. The phase of the cell cycle that matches the cyclin profile during *O. tsutsugamushi* infection is denoted by a red dashed box. **(f-k)** EA.hy926 cells were serum starved for 24 h before *O. tsutsugamushi* infection. At time of infection, serum-free medium was replaced with complete medium. WCLs were collected at 8, 24, 48, and 72 h p.i. and subjected to **(f)** western blot followed by densitometric analysis of **(g)** Cyclin D1, **(h)** Cyclin D3, **(i)** Cyclin E, **(j)** Cyclin A, or **(k)** Cyclin B1. Red asterisks (*) denote TSA56 in infected samples. The ratio of the densitometric signal for each cyclin relative to GAPDH was compared between infected and uninfected samples to determine the fold change at each timepoint. **(l-n)** Pharmacologic or siRNA-mediated inhibition of the cell cycle. **(l)** Experimental design employed for treatment of uninfected and infected host cells with cell cycle inhibitors. **(m)** EA.hy926 cells were treated with VEH (DMSO), medium without FBS (serum starved; SS), hydroxyurea (HU; 80 µM), or RO-3306 (9 µM; green) 24 h before infection. Treatment was maintained (closed circle) or removed at 48 h p.i. (open circle). Lysates were collected at 96 h p.i. **(n)** EA.hy926 cells were transfected with siNT, siCCNE1, siGMNN, or siCDC25a before being infected with *O. tsutsugamushi.* Lysates were collected at 72 h p.i. **(m-n)** *O. tsutsugamushi* GE mL^-1^ was determined by qPCR. Data are means ± SEM (*n* = 3-5 independent experiments). Schematics were generated using BioRender.com. Statistical significance was evaluated by One-way ANOVA followed by Tukey’s *post hoc* test. Statistically significant values (P < 0.05) are displayed.

To determine if S phase benefits *O. tsutsugamushi* fitness, we employed serum starvation to restrict the cell cycle in G0/G1^78^, hydroxyurea to arrest in S phase^79^, or RO-3306 to halt at the G2/M junction (Fig. 4l)^80^. The *Orientia* DNA load was measured at 96 h p.i. Each treatment’s effect on host cell metabolic activity was verified using the MTT (3-[4,5-dimethylthiazol-2-yl]-2,5-diphenyl-2H-tetrazolium bromide) assay^81^. Serum starvation and hydroxyurea had no effect whereas RO-3306 reduced metabolic activity by half (Supplementary Fig. 7d), the latter of which is expected during G2/M phase restriction^82^. Exposure of host cell-free *O. tsutsugamushi* to hydroxyurea and RO-3306 prior to infection did not impact its subsequent intracellular growth (Supplementary Fig. 7e). Whereas serum starvation and RO-3306 significantly lowered the bacterial yield, *Orientia* grew just as well in hydroxyurea treated cells as in controls (Fig. 4m). Treatment removal at 48 h p.i. to release cell cycle phase restriction did not bolster bacterial growth. As a complementary approach, cells were treated with siRNA targeting cyclin E to induce G1 arrest, geminin to induce S phase arrest, or cell division cycle 25A to induce G2 arrest^83^. Knockdown was confirmed at the time of infection (Supplementary Fig. 7f-k). Cells arrested in G1 and G2/M phase yielded significantly less *O. tsutsugamushi*, while the loads in cells arrested in S phase and control cells were comparable (Fig. 4n). Together, these results show that *O. tsutsugamushi* pauses the cell cycle at S phase to maximize its growth. Although the lack of p53 would facilitate transition to S phase, the delay in host cell proliferation observed for infected cells suggests that *Orientia* employs a p53-independent mechanism to arrest the cell cycle.

### *O. tsutsugamushi* inhibits host cell DNA damage

p53 is induced by DNA damage and is a key regulator of the DNA damage repair response^84^. Phosphorylation of histone H2.AX is a hallmark of DNA damage^85,86^. Phosphorylated H2.AX (ƔH2.AX) levels are unaltered in *O. tsutsugamushi* infected EA.hy926 cells until 72 h p.i. when they are significantly elevated (Fig. 5a, b). To determine if *O. tsutsugamushi* inhibits DNA damage, cells were treated with etoposide, which induces irreversible double stranded DNA breaks^87–89^. Treatment was initiated 24 h before each collection timepoint. Etoposide did not affect *O. tsutsugamushi* growth (Fig. 5c). As expected, etoposide strongly raised p53 and ƔH2.AX levels in uninfected cells (Fig. 5d-f). In infected cells, p53 levels were decreased even in the presence etoposide at all time points. ƔH2.AX levels were also reduced until 72 h p.i. at which time they had risen comparably to those in uninfected cells. Thus, *O. tsutsugamushi* inhibits DNA damage until late infection.

**Fig. 5.**
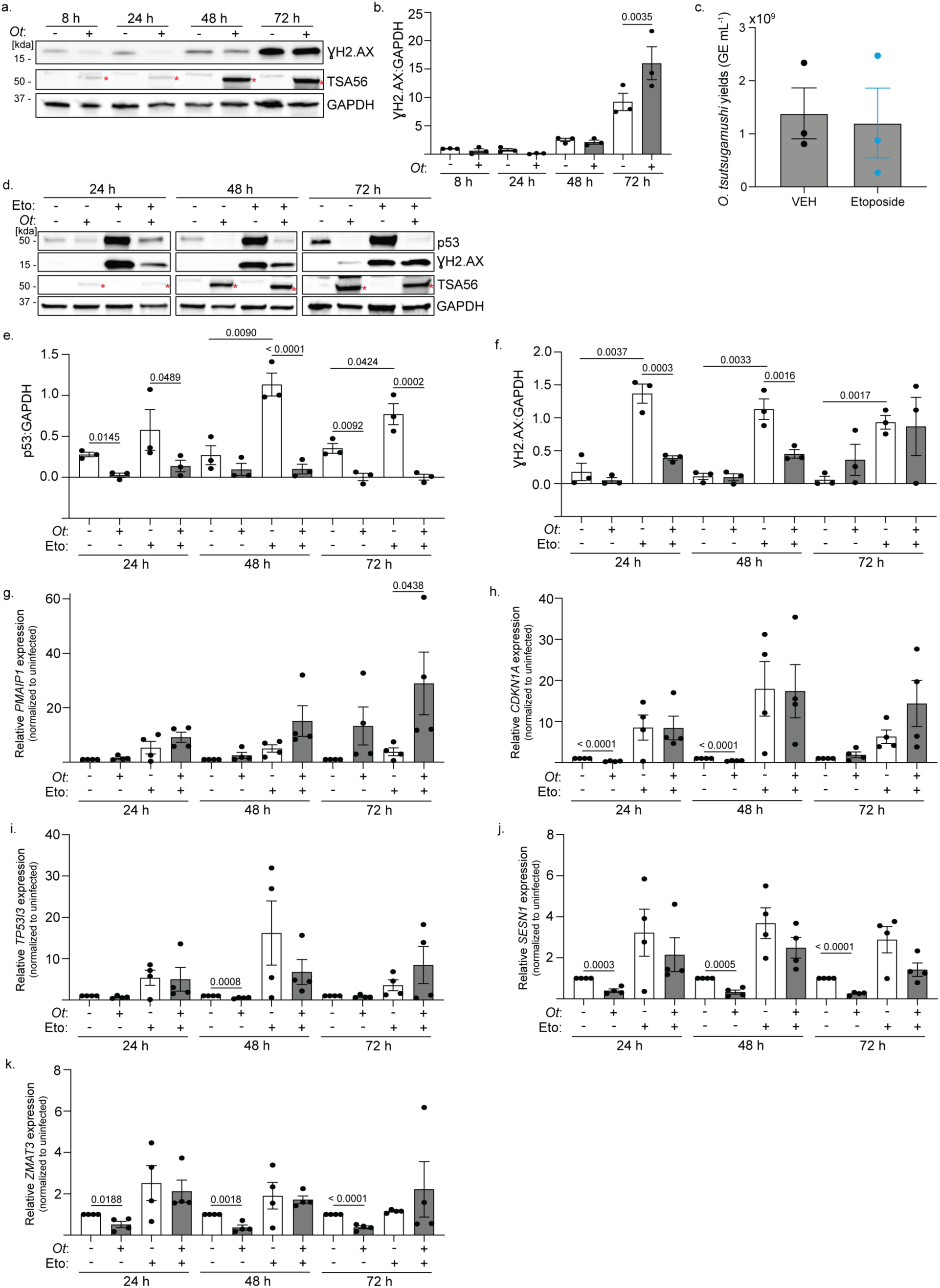
*O. tsutsugamushi* inhibits host cell DNA damage. **(a-b)** WCLs were collected from uninfected and *O. tsutsugamushi* infected EA.hy926 cells at 8, 24, 48, and 72 h p.i. followed by **(a)** western blot. Blots were probed with ƔH2.AX, TSA56, and GAPDH. The mean normalized ratio ± SEM of **(b)** ƔH2.AX:GAPDH was determined by densitometry analysis (*n* = 3 independent experiments). **(c-k)** EA.hy926 cells were infected and treated with etoposide (Eto; 1 µM) or vehicle (VEH) 24 h prior to collection. **(c)** *O. tsutsugamushi* GE mL^-1^ was determined using qPCR. Data are means ± SEM (*n* = 3 independent experiments). **(d-f)** WCLs collected at 24, 48, and 72 h p.i. were subjected to **(d)** western blot and **(f)** densitometry analysis. Blots were probed with p53, ƔH2.AX, TSA56, and GAPDH. **(g-j)** RT-qPCR was performed on total RNA collected at 24, 48, and 72 h p.i. Mean normalized ratios ± SEM of **(e)** p53:GAPDH or **(f)** ƔH2.AX:GAPDH, and relative gene expression of **(g)** *PMAIP1*, **(h)** *CDKN1A*, **(i)** *TP53I3*, **(j)** *SESN1*, and **(k)** *ZMAT3* normalized to *GAPDH* were calculated (*n* = 3-4 independent experiments). Red asterisks (*) denote TSA56 in infected samples. Statistical significance was evaluated by One-way ANOVA followed by Tukey’s *post hoc* test. Statistically significant values (P < 0.05) are displayed.

DNA damage stabilizes p53 which then activates a subset of DNA repair genes that promote senescence or cell death^90–92^. To further elucidate the effects that *O. tsutsugamushi*-driven p53 downregulation has on the DNA damage repair response, we treated cells with etoposide or vehicle and used RT-qPCR to measure expression of *PMAIP1* (phorbol-12-myristate-13-acetate-induced protein 1), *CDKN1A* (cyclin dependent kinase inhibitor 1A), *TP53I3* (tumor protein p53 inducible protein 3), *SESN1* (sestrin 1), and *ZMAT3* (zinc finger matrin-type 3). In vehicle treated infected cells, *CDKN1A*, *TP53I3*, *SESN1*, and *ZMAT3* were downregulated at all timepoints while *PMAIP1* was upregulated at 72 h p.i. (Fig. 5g-k). In response to etoposide, *PMAIP1* expression was more robustly induced in infected cells while *CDKN1A*, *TP53I3*, *SESN1*, and *ZMAT3* transcription increased similarly in uninfected and infected cells. Upregulation of *CDKN1A*, which encodes p21, during treatment with etoposide translated to an increase in p21 levels (Supplementary Fig. 8). Therefore, although *O. tsutsugamushi* does not induce expression of any of the assayed DNA damage response genes until late infection, it does not repress these genes in response to etoposide-induced genotoxicity.

### *O. tsutsugamushi* inhibits endothelial cell apoptosis

Because a major p53 function is to induce apoptosis^68^, we examined if *O. tsutsugamushi* infected endothelial cells are recalcitrant to this cell death pathway. Cleaved caspase 3, caspase 7, and PARP1, which are apoptosis hallmarks^93^, were detected in uninfected cells exposed to etoposide but not in vehicle-or etoposide-treated infected cells (Fig. 6a-d). Staurosporine is a pan-kinase inhibitor that induces apoptosis by p53-dependent and -independent mechanisms that do not directly induce DNA damage^94–96^. *O. tsutsugamushi* suppressed p53 levels and caspase 3, caspase 7, and PARP1 cleavage in staurosporine treated cells (Supplementary Fig. 9). To account for caspase-independent mechanisms of apoptosis/cell death, we used flow cytometry to assay for annexin V binding to phosphatidylserine on the plasma membrane and uptake of membrane-impermeable propidium iodine (PI). In the absence of infection, etoposide and staurosporine robustly increased numbers of apoptotic/dead cells (Fig. 6e, f). This effect was not observed for infected cells until 72 h p.i. Thus, *O. tsutsugamushi* inhibits both DNA damage-dependent and - independent apoptosis of endothelial cells until late infection.

**Fig. 6.**
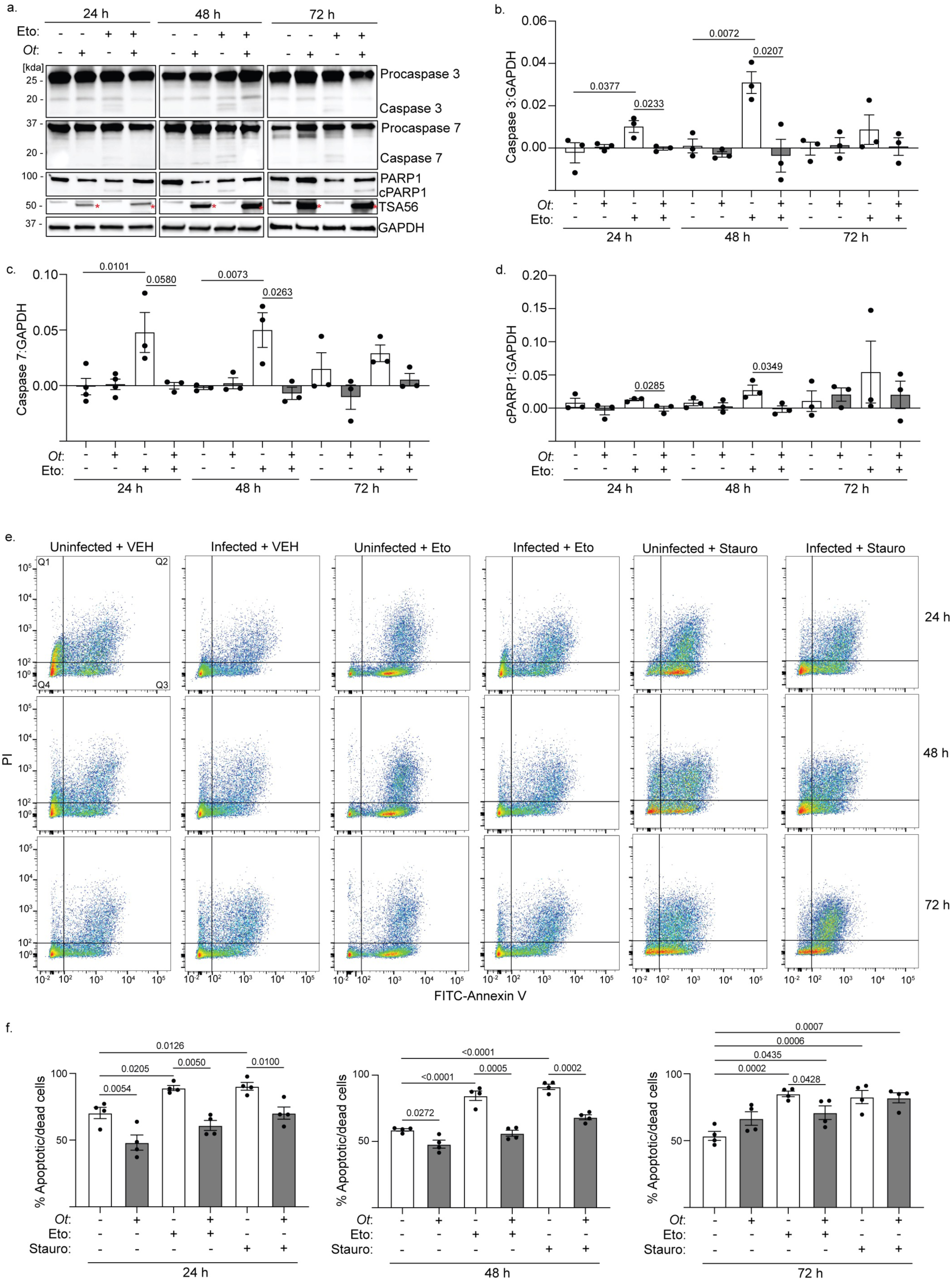
*O. tsutsugamushi* inhibits endothelial cell apoptosis. **(a-d)** Uninfected and infected EA.hy926 were treated with etoposide (Eto) 24 h prior to collection. **(a)** Western blots of WCLs collected at 24, 48, and 72 h p.i. were probed for procaspase 3 and cleaved caspase 3, procaspase 7 and cleaved caspase 7, full length and cleaved (cPARP1) PARP1, TSA56, and GAPDH. Red asterisks (*) denote TSA56 in infected samples. Mean normalized ratios ± SEM of **(b)** Caspase3:GAPDH, **(c)** Caspase7:GAPDH, and **(d)** cPARP1:GAPDH densitometric signals were calculated (*n* = 3 independent experiments). **(e-f)** VEH, staurosporine (Stauro)-treated, or Eto-treated uninfected and infected samples were collected at 24, 48, and 72 h p.i. and stained with FITC-Annexin V and propidium iodine (PI). **(e)** Samples were subjected to flow cytometry. Quadrants displayed on dot plot of uninfected + VEH-treated sample represent the quadrants across all samples. Quadrant 1 (Q1; FITC-/PI+) and Q2 (FITC+/PI+) contain late apoptotic and dead cells, respectively. Q3 (FITC+/PI-) consists of early apoptotic cells and Q4 (FITC-/PI-) contains live cells. **(f)** The percentage of apoptotic cells/dead (Q1-3) was determined. The mean percentage of cells ± SEM of each quadrant was calculated (*n* = 3 independent experiments). Statistical significance was evaluated by One-way ANOVA followed by Tukey’s *post hoc* test. Statistically significant values (P ≤ 0.05) are displayed.

### *Orientia* Ank13 downregulates *TP53* to inhibit DNA damage-induced apoptosis

We rationalized that the chloramphenicol-sensitive ability of *O. tsutsugamushi* to downregulate *TP53* involves a bacterial protein that modulates host cell gene expression. *Orientia* encodes one of the largest cadres of ankyrin repeat (AR)-containing effectors (Anks), most of which exhibit a bipartite architecture consisting of an N-terminal AR domain and a C-terminal F-box^56,97–100^. ARs mediate protein-protein interactions, and pathogen acquisition of genes encoding AR proteins is linked to increased virulence^99,101,102^. The F-box is a eukaryotic-like motif that coopts the host SCF E3 ubiquitin ligase complex and is essential for multiple *Orientia* Anks to modulate host cell processes^49–51,98,103^. Ank13 is an *O. tsutsugamushi* nucleomodulin first documented to be expressed during infection of HeLa cells^49^. It has eight ARs and an F-box (Supplementary Fig. 10a). Its nucleotropism depends on I127 located in AR4. When ectopically expressed in HeLa cells, Ank13 dysregulates expression of more than 2,000 genes in nucleotropism- and F-box-dependent manners^49^. We confirmed that *O. tsutsugamushi* expresses *ank13* throughout infection of EA.hy926 cells (Supplementary Fig. 10b, c). In the absence of tools for genetically manipulating *O. tsutsugamushi*^104^, we determined if Ank13 can downregulate *TP53* and p53 expression by transfecting EA.hy926 cells to express GFP-Ank13, nucleotropism-defective GFP-Ank13_I127R49_, or GFP. Cells were also transfected to express GFP-Ank13-F-boxAAAAA, which cannot bind the SCF complex due to its five F-box residues essential for doing so being replaced with alanine^49,98,105^. GFP-positive cells were enriched for by fluorescent-activated cell sorting followed by RT-qPCR or immunoblot analysis. Cells expressing GFP-tagged Ank13 and Ank13-F-boxAAAAA but not Ank13_I127R_ had significantly lower *TP53* transcript along with reduced p53 and ƔH2.AX protein levels compared to cells expressing GFP (Fig. 7a-d). Moreover, GFP-Ank13 and GFP-Ank13-F-boxAAAAA but not GFP-Ank13_I127R_ inhibited expression of the *TP53* P1 isoform (Fig. 7e), while none altered *TP53* P2 isoform expression (Fig. 7f). Thus, ectopically expressed Ank13 phenocopies *O. tsutsugamushi* infection by specifically inhibiting P1 *TP53* isoform expression in a nucleotropism-dependent but F-box-independent manner to reduce p53 levels. The nucleotropism of Ank13 is also key for it to protect endothelial cells from DNA damage. Finally, GFP-Ank13 prevented endothelial cell apoptosis induced by etoposide but not staurosporine and required both its nucleotropism and F-box to do so (Fig. 7g, h). Overall, Ank13 is linked to the ability of *O. tsutsugamushi* to impede p53 expression, protect against genotoxicity, and inhibit DNA damage-dependent but not -independent apoptosis.

**Fig. 7.**
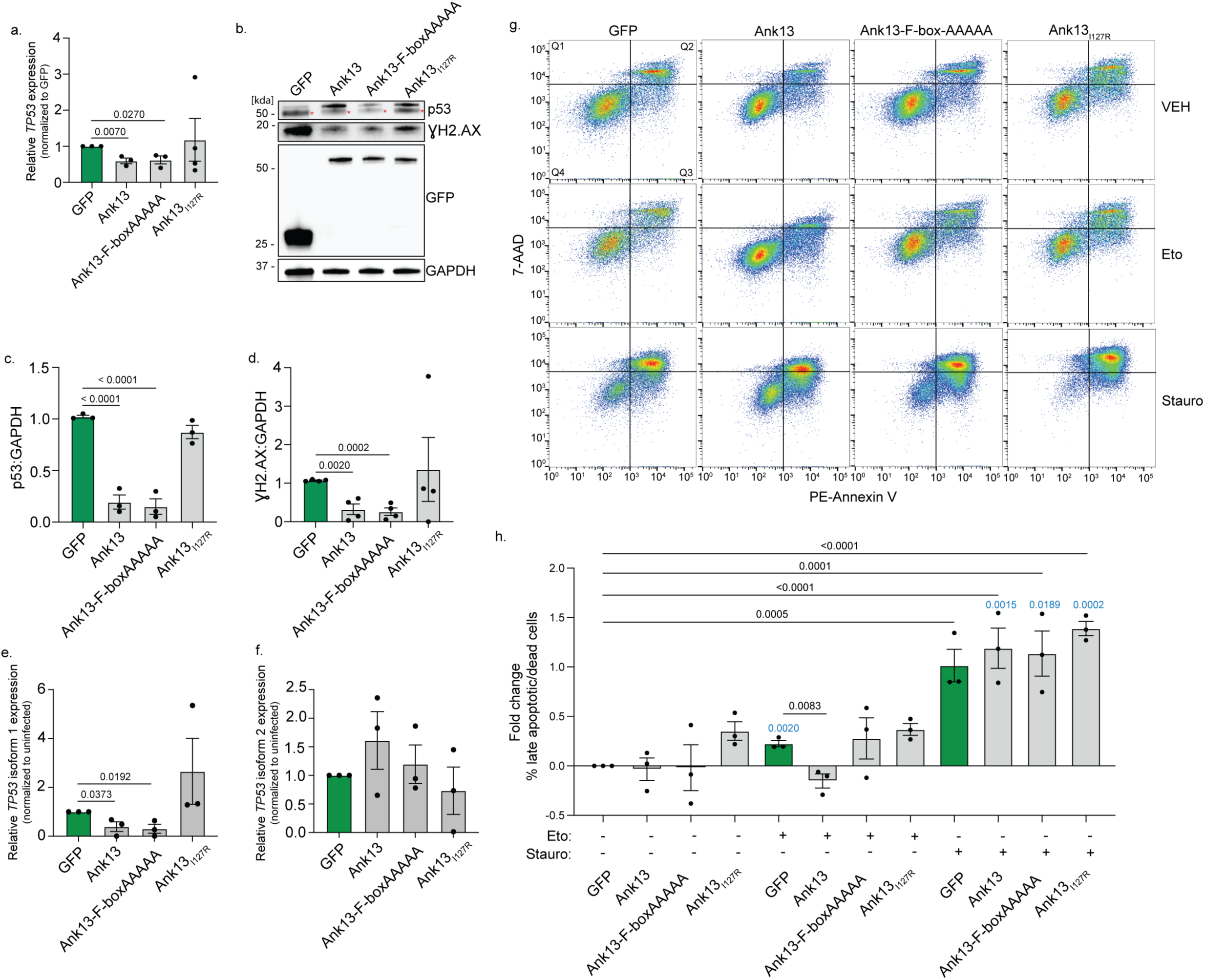
Ank13 downregulates *TP53* to inhibit DNA damage-dependent apoptosis. **(a-f)** RNA and WCLs were collected from fluorescence-activated cell-sorted EA.hy926 cells expressing GFP, GFP-Ank13, GFP-Ank13ΔF-boxAAAAA, or GFP-Ank13_I127R_ and subjected to **(a, e, f)** RT-qPCR and **(b)** western blot, respectively. Blots were probed for p53, ƔH2.AX, GFP, and GAPDH. Red asterisks (*) denote p53. Mean normalized ratios ± SEM of **(a)** *TP53*, **(e)** *TP53* P1, and **(f)** *TP53* P2 to *GAPDH* and **(c)** p53:GAPDH and **(d)** ƔH2.AX:GAPDH densitometric signal ratios were calculated (*n* = 3-4 independent experiments). **(g-h)** EA.hy926 cells transfected to express GFP or the indicated GFP-tagged Ank13 proteins were treated with Stauro, Eto, or VEH and stained with PE-Annexin V and 7-AAD. **(g)** Samples were subjected to flow cytometry. Quadrants displayed on the dot plot of GFP + VEH represent the quadrants for all samples. Quadrant 1 (Q1; FITC-/PI+) and Q2 (FITC+/PI+) consists of late apoptotic and dead cells, respectively. Q3 (FITC+/PI-) contains early apoptotic cells.Q4 (FITC-/PI-) contains live cells. **(h)** The percentage of apoptotic cells/dead cells (Q1-3) was determined. The fold change in the mean percentage of cells ± SEM were calculated (*n* = 3 independent experiments). Statistical significance was evaluated by One-way ANOVA followed by Tukey’s or Dunnett’s *post hoc* test for comparison of each sample to each other (black) or each sample to the VEH-treated counterpart (blue), respectively. Statistically significant values (P ≤ 0.05) are displayed.

## DISCUSSION

Apoptosis is a key innate defense against intracellular microbes that eliminates the replicative niche and is elicited in response to DNA damage incurred during infection. *O. tsutsugamushi* is the first pathogen identified to delay DNA damage-dependent apoptosis by blocking *TP53* expression. *Orientia* couples this strategy with protecting its endothelial host cell DNA from insult in a p53-independent manner and modulating the cell cycle to yield an intracellular environment that favors its logarithmic growth. Many pathogens that target p53 do so by directing its proteasomal degradation^5–13,28^. Yet, as demonstrated for *Chlamydia trachomatis* and *Plasmodium yoelii*, this strategy is an Achille’s heel that can be nullified using Nutlin 3a as a host-directed therapeutic to restore p53 levels and reduce the pathogen load^8,12^. Because the *Orientia* mechanism is HDM2- and proteasome independent, Nutlin 3a would be ineffective against scrub typhus.

The abilities of *O. tsutsugamushi* to inhibit *TP53* expression, protect against host cell genotoxicity, and delay apoptosis are linked, at least in part, to Ank13, as it singularly executes all three promicrobial strategies when ectopically expressed. *O. tsutsugamushi*-mediated p53 reduction is first observed during the initial hours of infection, which agrees with its high turnover rate and implies the action of a preloaded effector like Ank13. Indeed, because *Orientia* expresses Ank13 throughout its infection cycle, it would be ready for deployment upon invasion. Ank13 blocks expression of the P1 but not P2 *TP53* isoform to recapitulate how *Orientia* targets the *TP53* upstream region to deplete host cells of p53. The effector does so in a nucleotropism-dependent manner, indicating that it acts in the nucleus to block *TP53* at the transcriptional level by binding DNA at the *TP53* promoter region, mRNA at the 5’-untranslated region, or a transcription factor. For the latter possibility, because Ank13 *TP53* inhibition is F-box-independent, it would impair the target transcription factor without ubiquitinating or directing it for proteasomal degradation. The mechanism(s) by which nucleotropic Ank13 impedes *TP53* expression are likely also employed to alter expression of more than 2,000 genes^49^.

*O. tsutsugamushi* inhibits apoptosis using Ank13-dependent and -independent strategies. Ank13 protects against DNA damage responses induced by cationic lipid transfection or etoposide in a nucleotropism-dependent manner, which phenocopies *Orientia* prevention of double-stranded DNA breaks during early infection^52,55^. The ability of Ank13 to impair etoposide-induced apoptosis requires both its nucleotropism- and F-box, the latter of which echoes how *O. tsutsugamushi* Ank5 uses its F-box to orchestrate ubiquitination and proteasomal degradation of NLRC-5 (NOD-like receptor family CARD domain containing 5) to prevent major histocompatibility class I expression^50^. Ank13 may promote degradation of one or more host factors that drive apoptosis due to irreparable DNA damage. Ank13 is not involved in the resistance to staurosporine-induced apoptosis exhibited by *Orientia* infected cells^51^, reinforcing that it counters apoptosis exclusively in a genoprotective manner. Rather, resistance to apoptosis elicited by staurosporine is conferred by a subset of five F-box-containing Anks (Ank1, Ank5, Ank6, Ank9, Ank17) that retain Cul1 in the cytoplasm to lower the Cul1:c-MYC intranuclear balance and thereby dysregulate expression of c-MYC target genes involved in apoptosis^51^. Thus, at least six *Orientia* Anks transcriptionally inhibit apoptosis. *O. tsutsugamushi* also exerts an anti-apoptotic effect by impeding intracellular calcium release via an unknown mechanism^106^.

*O. tsutsugamushi* reduction of p53 levels promotes an endothelial cell transcriptional program that would normally culminate in uncontrolled cell cycle progression and proliferation. Yet, the pathogen modulates this response by arresting the cell cycle at S phase to promote its logarithmic growth. How might S phase arrest benefit *Orientia*? Following invasion, it traffics to the centrosome where it replicates. The resulting bacterial population remains there until exiting to initiate the next infection wave^42^. Because the centrosome duplicates during G2 and the duplicated centrosomes separate during the G2/M transition^107^, perhaps *Orientia* orchestrates the pause at S phase to prevent disruption of its preferred intracellular location for growth. Also, cells in S phase undergo metabolic shifts to support DNA replication and cell division, becoming enriched in amino acids and other intermediates that can feed into the tricarboxylic acid cycle (TCA)^108^. Conspicuously, *O. tsutsugamushi* growth corresponds with reductions in both amino acids and TCA intermediates^53^. Moreover, p53 exerts antimicrobial activity by inhibiting glycolysis and the pentose phosphate phosphate pathway, both of which feed into the TCA^2,108^. By simultaneously downregulating *TP53* and arresting the cell cycle at S phase, *Orientia* may synergistically enrich its intracellular environment with the host metabolites that fuel its growth.

The majority of endothelial cell genes upregulated during *O. tsutsugamushi* replication include interferon, NF-κB, and IL-10 signaling as well as inflammasome activation. This indicates that antimicrobial responses, many of which are classically defined as antiviral, are mounted at the transcriptional level in response to late-stage *Orientia* infection and is consistent with observations for scrub typhus patient and animal model studies^109–114^. However, p53 contributes to interferon-mediated innate immunity, and the p53 downstream target, p21, which is also depleted during *Orientia* infection, is an important interferon-induced antiviral restriction factor^4,112^. By blocking *TP53* expression, *O. tsutsugamushi* at least partially counters these responses. Furthermore, even though NF-κB responses are transcriptionally upregulated in *Orientia* infected cells and loss of p53 can lead to increased NF-κB activity^115^, the bacterium inhibits NF-κB-dependent gene expression by impairing the transcription factor’s accumulation in the nucleus^103^. This strategy minimally involves the actions of *Orientia* Ank1 and Ank6 and bacterial stabilization of the host NF-κB inhibitor, p105^103,116^. Interestingly, the only group of host transcripts downregulated when *Orientia* load is high are involved in translation elongation. Many intracellular pathogens counteract host translation to prevent proinflammatory protein production^117–120^. Previous work showed that *O. tsutsugamushi* Boryong strain effectors TPR43 and TRP46 inhibit eukaryotic translation elongation^121^. Also, our lab demonstrated that Ikeda Ank4 and Ank9 disrupt the unfolded protein response^122,123^, which is directly linked to an inhibition of host translation^117,118^. Therefore, inhibiting host cell translation might be yet another means by which *O. tsutsugamushi* facilitates its intracellular survival.

Downregulating *TP53* to deplete p53 is a strategy that *O. tsutsugamushi* Ikeda shares with UT76, Karp, and Gilliam. The complement of Ank genes varies among all sequenced isolates, and of the four examined herein, only Ikeda and UT76 encode Ank13^97^. Thus, *O. tsutsugamushi* strains utilize Ank13 and unidentified virulence factors to block *TP53* expression. Regardless, the overall conservation of this tactic underscores its importance to *Orientia* fitness and suggests that p53 suppression during scrub typhus is common. Although largely considered an acute illness^31,41,42^, limited studies have suggested that scrub typhus can become latent^124,125^. While cells heavily infected with *O. tsutsugamushi* that have accumulated DNA damage would presumably be eliminated by apoptosis during acute disease, the intracellular bacterial burden and whether unrepaired genotoxicity is incurred during latent infection are unknown. Given that infectious diseases cause up to 20% of human cancers and *O. tsutsugamushi* infection can elevate the risk for other serious health conditions later in life^1,2,126^, it might be worthwhile for longitudinal studies to assess the epidemiological relationship between scrub typhus and subsequent development of cancer.

In summary, *O. tsutsugamushi* is the first example of a pathogen that lowers p53 host cell levels by downregulating *TP53* expression. In the absence of p53, which would be expected to lead to unchecked genotoxicity and cell cycle progression, *Orientia* modulates both. During its logarithmic replication, it protects against DNA damage, inhibits DNA damage-induced apoptosis, and mediates cell cycle arrest at S phase, the latter of which likely affords the bacterium needed metabolites. The abilities of *Orientia* to block *TP53* expression, mediate genoprotection, and impair DNA damage-dependent apoptosis involve its nucleotropic effector, Ank13. This study furthers understanding of how *O. tsutsugamushi* and potentially other intracellular pathogens modulate p53 and p53-related pathways to maintain their niches for as long as possible.

## METHODS

### Cultivation of cell lines and *O. tsutsugamushi* infections

Uninfected HeLa human cervical epithelial cells (CCL-2; American Type Culture Collection [ATCC], Manassas, VA) and EA.hy926 human umbilical vein cells (CRL-2922; ATCC) were maintained as done previously. HeLa cells infected with *O. tsutsugamushi* strains Ikeda, UT76, Karp, or Gilliam were maintained as described^50,53^. To isolate *O. tsutsugamushi* from infected host cells, media was removed and the cells were washed with 1X phosphate-buffered saline (PBS; 1.05 mM KH_2_PO_4_, 155 mM NaCl, 2.96 mM Na_2_HPO_4_ [pH 7.4]) before trypsinization. Dislodged cells were pelleted at 1000 x*g* for 3 min, the supernatant removed, and the pellet resuspended in 1X PBS. The resuspended pellet was combined with 0.7 mm zirconia beads (Biospec Products; Bartlesville, OK), and the host cells were disrupted to release intracellular bacteria using an analog disruptor genie (Scientific Industries, Inc.; Bohemia, New York) for 2 min. Beads and host cell debris were pelleted at 1000 x*g* for 3 min. The supernatant containing host cell-free *O. tsutsugamushi* was collected and utilized for continuous cultivation in HeLa cells or for infection studies in EA.hy926 cells^53^. Synchronous infections were achieved by removing inocula and replacing with fresh complete Dulbecco’s modified Eagle’s medium (DMEM) with l-glutamine, 4.5 g d-glucose and 100 mg sodium pyruvate (Gibco, Gaithersburg, MD) supplemented with 10% (vol/vol) heat-inactivated fetal bovine serum (FBS) (Gemini Bioproducts, West Sacramento, CA), 1X minimal essential medium containing non-essential amino acids (Gibco) and 15 mM HEPES (Gibco) at 2 to 4 h p.i. Experiments were verified for achieving a multiplicity of infection (MOI) of 5-10 in at least 75% of the host cells by assessing coverslips of infected cells by immunofluorescence as described below. Samples in which less than 75% of the cells were infected were not used.

### Genome equivalents (GE) quantification

Lysates for GE were collected by trypsinization followed by dilution in H_2_O and incubation at 100°C for 15 min as described previously^127^. Briefly, at the time of analysis, samples were thawed, vortexed, and GE quantified by qPCR. *O. tsutsugamushi* str. Ikeda *tsa56* gene (OTT_RS04590)^53^ or *tsa47* (47-kDa type-specific antigen) gene (OTT_ RS06370) were detected using gene-specific primers (Supplementary Table 1) with PerfeCTa SYBR Green Fastmix (Quantbio, Beverly, MA), and a CFX384 Real Time PCR Detection System (Bio-Rad Laboratories). Thermal cycling conditions were 95°C for 30s followed by 40 cycles of 95°C for 10s to 54°C for 10s, and a 65°C-to-95°C melt curve. A standard curve was prepared using pCR4.0::*Ot_tsa56*^50^ or pCR4.0::*Ot_tsa47*. pCR4.0::*Ot_tsa47* standard plasmid was generated by Azenta Genewiz from Azenta Life Sciences (https://www.genewiz.com/en/Public/Services/Gene-Synthesis/Gene-Synthesis) to include base pairs 354-844 of the *tsa47* gene. *O. tsutsugamushi* GE mL^-1^ was calculated based on the standard curve as done previously^53,127^.

### RNA-seq

EA.hy926 cells at 100% confluency were synchronously infected with *O. tsutsugamushi*. At 8, 48, and 72 h p.i., spent medium was removed and the cells washed with 1X PBS before trypsinization. Dislodged cells were pelleted at 1000 x*g* for 3 min, and total RNA was extracted using RNeasy mini kit (Qiagen [74104], Hilden, Germany). GE per sample were measured per timepoint. RNA concentration and purity were assessed using a NanoVue Plus Spectrophotometer (GE HealthCare, Chicago, IL) before storage at -80°C. Three independent biological samples that met the requisite criteria of ≥ 20 ng/uL, OD_260/280_ ≥ 2.0, and OD_260/230_ ≥ 2.0 were submitted to Novogene (Sacramento, CA) where sample integrity was confirmed using an Agilent 2100 Bioanalyzer (Agilent Technologies, Santa Clara, CA) prior to sequencing. Samples with an RNA integrity number of ≥ 4.0 and no gDNA contamination were sequenced on an Illumina high-throughput sequencing platform (Illumina, San Diego, CA). Spliced Transcripts Alignment to a Reference (STAR) program was employed to map raw reads to human reference genome (GRCh38/hg38)^128^. DESeq2 R software package 1.14.1 (Bioconducter, https://www.bioconductor.org/) was used to perform differential expression significant analysis between infected and uninfected samples per timepoint^129^. To control the false discovery rate (FDR) of the resulting P values, the Benjamini and Hochberg’s approach was employed and adjusted P values generated^130,131^. Genes with an adjusted P value of ≤ 0.05, as well as a log_2_(fold change) > 1 or < -1 of significantly differentially expressed genes in infected cells compared to the uninfected controls were considered differentially expressed genes (DEGs). DESeq2 R package was used to determine the Pearson’s correlation coefficient and PCAtools R package (Bioconducter) was employed to perform a principal components analysis.

### STRING analysis

Interaction networks were defined using STRING^132^. Briefly, DEGs per timepoint were individually imported into STRING, and unconnected nodes were removed. MCL clustering was performed with a constricting inflation parameter of 2. GO terms sub-grouped into biological processes per cluster were defined and those with a FDR ≤ 0.05 were considered significantly enriched pathways.

### qRT-PCR

Total RNA was isolated from uninfected or *O. tsutsugamushi* infected EA.hy926 cells, and concentration and purity were determined as described above. One µg of RNA was treated with amplification grade DNase I (Invitrogen) following the manufacturer’s protocol. Next, iScript Reverse Transcription Supermix (Bio-Rad) was used to generate cDNA. To ensure gDNA was successfully removed, reactions in the absence or presence of reverse transcriptase were subjected to PCR with MyTaq polymerase (Bioline, Taunton, MA) and primers for human *GAPDH*^133^ before being visualized by an agarose gel electrophoresis. gDNA-free cDNA was subjected to qPCR with PerfeCTa SYBR Green Fastmix and primers for *TP53, HDM2, TP53* P1, *TP53* P2, *PMAIP1, CDKN1A, TP53I3, SESN1, ZMAT3* (Supplementary Table 1)*, GAPDH*^133^*, O. tsutsugamushi* 16S rDNA (*ott16s*)^133^, or *ank13*^49^. Thermal cycling conditions used were 95 °C for 30 s, followed by 40 cycles of 95 °C for 5 s and 55 °C for 5 s, followed by a melt curve from 60 °C to 95 °C. 2^-ΔΔCT^ was calculated to determine the relative expression of each gene^50^.

### Western blotting

Cells were washed with 1X PBS, harvested by trypsinization, pelleted at 1,000x*g* for 3 min, and resuspended in radioimmunoprecipitation assay (RIPA) buffer (50 mM Tris-HCl [pH 7.4], 150 mM NaCl, 1% NP-40, 1% sodium deoxycholate, 1 mM EDTA [pH 8]) containing Halt protease and phosphatase inhibitor cocktail (ThermoFisher Scientific, Waltham, MA). Suspensions were incubated on ice for 15-45 min and insoluble fractions were pelleted by centrifugation at 16,000 x*g* for 10 min. Supernatants were collected and stored at -20°C. Protein concentration was quantified using the Bradford assay (Bio-Rad [#5000006], Hercules, CA). Equivalent amounts of protein were resolved on a 4 to 15% TGX polyacrylamide gels (Bio-Rad) at 110 V for 15 min followed by 200 V from 30 min. Proteins were transferred onto nitrocellulose membrane (Bio-rad; #1620115) in Towbin buffer (24mM Tris, 192 mM glycine, 20% [vol/vol] methanol) at 100 V for 30 min. Blots were blocked with either 5% (vol/vol) nonfat dry milk or 5% (vol/vol) BSA in Tris-buffered saline plus Tween-20 (TBS-T; 25 mM Tris HCl, 137mM NaCl, 2.7 mM KCl, 0.05% Tween-20, pH 7.4) for 30-60 min before incubation with primary antibody. Primary antibodies used were: mouse anti-p53 (Santa Cruz [sc-98]; 1:200), rabbit anti-p21 (Cell Signaling [#2947]; 1:1,000), mouse anti-HDM2 (Invitrogen [#33-7100]; 1:500), mouse anti-Cyclin E (Santa Cruz, Dallas, TX [sc-377100]; 1:500), mouse anti-Cyclin A (Santa Cruz [sc-271682]; 1:250), rabbit anti-Cyclin D3 (Cell Signaling, Danvers, MA [#2936]; 1:1,000), rabbit anti-Cyclin D1 (Cell Signaling [#55506]; 1:1,000), rabbit anti-Cyclin B1 (Cell Signaling [#4138]; 1:1,000), rabbit anti-GFP (Invitrogen [A11122]; 1:1000), rabbit anti-GMNN (Cell Signaling [#52508]; 1:1,000), rabbit anti-cdc25a (Cell Signaling [#3652]; 1:1,000), rabbit anti-Caspase 7 (Cell Signaling [#9492]; 1:1,000), rabbit anti-Caspase 3 (Cell Signaling [#9662]; 1:1,000), rabbit anti-PARP1 (Invitrogen [MA5-15031]; 1:1,000), rabbit anti-PKR (Cell signaling [#3072; 1:1,000), rabbit anti-p-PKR (T446) (Abcam [ab32036]; 1:1,000), rabbit anti-*O. tsutsugamushi* TSA56^123^ (1:1,000), mouse anti-GAPDH (Santa Cruz [sc-365062]; 1:750). Horseradish peroxidase-conjugated horse anti-mouse or anti-rabbit (Cell Signaling Technology [mouse #7076, rabbit #7074]; 1:10,000) were used to detect bound primary antibodies. Blots were incubated with SuperSignal West Pico PLUS, SuperSignal West Dura, or SuperSignal West Femto chemiluminescent substrate (ThermoFisher Scientific) before visualization using the ChemiDoc Touch Imaging System (Bio-Rad). Densitometric values were determined using the Bio-Rad Image Lab 6.0 software.

### Immunofluorescent microscopy

EA.hy926 cells were seeded in 24-well plates with 12-mm round glass coverslips (Electron Microscopy Sciences, Hatfield, PA) and synchronously infected with *O. tsutsugamushi*. The cells were fixed with methanol for 5 min followed by successive incubations with rabbit-anti TSA56^123^ (1:1,000) for 1 h in 1X PBS containing 5% (vol/vol) BSA and Alexa Fluor 488-conjugated goat anti-rabbit IgG (Invitrogen [A-11034]; 1:1000) at 1:1,000 with the addition of 0.1 mg mL^-1^ 4′,6′-diamidino-2-phenylindole (DAPI; Invitrogen [D1306]). The coverslips were then mounted using ProLong Gold antifade reagent (Invitrogen) and imaged with a TCS SP8 microscope (Leica Microsystems, Germany).

### Pharmacologic treatments

Chloramphenicol (CAM; 34 ug/mL) was added to uninfected and *O. tsutsugamushi* infected EA.hy926 cells at 2 h p.i. EA.hy926 cells were treated with Nutlin 3a (5 or 10 µM)^6^, C16 (100 nM), (hydroxyurea (HU; 80 µM)^80^, or RO-3306 (9 µM)^134^ 18-24 h before infection with *O. tsutsugamushi*, and etoposide (1 µM)^87–89^ or staurosporine (1 µM)^94,135^ 18-24 h prior to collection. Pharmacologic agents were diluted in complete DMEM and remained in culture unless otherwise stated. The effects of each pharmacologic inhibitor and serum starvation on host cell fitness was assessed by the MTT assay as described^53^. To determine the effects of hydroxyurea and RO-3306 on host cell-free *O. tsutsugamushi*, the bacteria were treated 30 min prior to being added to untreated EA.hy926 cells followed by GE quantification^50,53,127^.

### Proliferation assay

EA.hy926 cells were seeded to achieve 60-70% confluency and serum starved in DMEM without FBS, HEPES, and non-essential amino acids for 24 h. After serum starvation, cells were labeled with the CellTrace Far Red Proliferation kit (ThermoFisher Scientific [C34564]) following the manufacturer’s protocol. Briefly, the medium was removed, 1 µM of CellTrace Far Red dye diluted in pre-warmed 1X PBS was added, and the cells were incubated at 35°C in a humidified incuabor with 5% CO_2_ for 20 min. The dye solution was removed, and cells were washed twice with complete DMEM. Cells were incubated in complete DMEM at 35°C with 5% CO_2_ for 10 min prior to *O. tsutsugamushi* infection. At the time of analysis, cells were dislodged with 0.526 mM EDTA (Versene) solution and the fluorescence intensity of CellTrace Far Red dye in 10,000 individual cells was quantified using a Becton Dickinson (BD) FACSMelody with BDFACSChorus 1.3.3 software (BD Biosciences, Franklin Lakes, NJ). Analysis was performed in FlowJo (version 10.8.1) software (BD Biosciences) with population gates isolating single cell populations. In parallel, immunofluorescence microscopy was performed with the adjustment of serum starvation prior to infection, and percentage of infected host cells at each timepoint was verified^50^.

### RNAi

EA.hy926 cells grown to 100% confluency were transfected with ON-TARGETplus human Cyclin E, CDC25A, or GMNN SMARTpool siRNA, or non-targeting siRNA (GE Healthcare Dharmacon, Inc., Lafayette, CO) at a final concentration of 30 pmol following the manufacturer’s protocol. After 48 h, siRNA was replenished, and cells were infected with *O. tsutsugamushi*. GE was measured for uninfected and *O. tsutsugamushi* infected samples.

### Plasmid constructs

pGFP-Ank13 and GFP-Ank13_I127R_ were generated previously^49^. In-Fusion Cloning Primer Design Tool v1.0 (www.takarabio.com) was used to designs primers to generate GFP-Ank13-F-boxAAAAA (Supplementary Table 1), which contained the subsequent point mutations, I451A, P452A, E454A, I459A, and D467A. GFP-Ank13-F-boxAAAAA was generated following the TaKaRa Bio USA (San Francisco, CA) In-Fusion Mutagenesis protocol with pGFP-Ank13 as a template^49^.

### Transfections

EA.hy926 cells were grown to 100% confluency before transfection with GFP, GFP-Ank13, GFP-Ank13-F-boxAAAAA, or GFP-Ank13_I127R_ plasmid DNA using Lipofectamine 2000 (Invitrogen) and incubation at 37°C in a humidified incubator with 5% CO_2_. At 18-24 h post transfection, samples were washed with 1X PBS, trypsinized, and subjected to fluorescent-activated cell sorting (FACS) for collection of GFP positive cells as done previously^50^. Total RNA for RT-qPCR or lysates for immunoblot analysis were collected from enriched GFP populations.

### Annexin V staining

Uninfected or *O. tsutsugamushi*-infected cells, or cells expressing GFP, GFP-Ank13, GFP-Ank13ΔF-boxAAAAA, or GFP-Ank13_I127R_ were treated with vehicle, etoposide, or staurosporine for 24 h. At the time of collection, uninfected and infected samples were incubated with FITC-Annexin V and propidium iodine (PI), while cells expressing GFP or GFP-Ank13 and mutants were incubated with PE-Annexin and 7-AAD and analyzed by flow cytometry on a BD FACSMelody with BDFACSChorus 1.3.3 software (BD Biosciences) as described^51^. Compensation and quadrant analysis were performed using FlowJo software (version 10.8.1; BD Biosciences). Cells in quadrant 1 (Q1; PI+/FITC- or 7-AAD+/PE-) and Q2 (PI+/FITC+ or 7-AAD+/PE+) were considered late apoptotic/dead. Cells in Q3 (PI-/FITC+ or 7-AAD-/PE+) were deemed early apoptotic while those in Q4 (PI-/FITC- or 7-AAD-/PE-) were considered live. The fold change in the percentage of late apoptotic/dead cells was calculated by subtracting the percentage of late apoptotic/dead cells per each vehicle control-treated sample from that of its etoposide- or staurosporine-treated counterpart and dividing the difference by its respective vehicle control.

### Statistical analysis

Statistical analyses were performed using Prism 9.0 (GraphPad, San Diego, CA). Significance between two groups was determined by an unpaired student’s *t*-test. One-way analysis of variance followed by Tukey’s honest significance or Dunnett’s *post hoc* tests was used to test for significant differences between groups or each group compared to a control, respectively. Statistical significance was set at P values of ≤ 0.05.

## ACKNOWLEDGEMENTS

We thank Dr. Haley E. Adcox, Dr. Hanna J. Laukaitis-Yousey, Dr. E. Handly Mayton, and Jason R. Hunt for critical review of this manuscript. This work was supported by National Institutes of Health-National Institute of Allergy and Infectious Diseases (www.niaid.gov) grants 1R01 AI167857 to J.A.C. and 1F32 AI174656A1 to P.E.A.

## AUTHOR CONTRIBUTIONS

Conceptualization, P.E.A. and J.A.C.; investigation, P.E.A., D.L.A., S.B..; writing – original draft, P.E.A. and J.A.C.; writing – review and editing, P.E.A. and J.A.C.; supervision, J.A.C.; acquisition of grant support, J.A.C. and P.E.A.

