## Supplementary Table 1, Supplementary Figures 1-10 for "Obligate intracellular *Orientia tsutsugamushi* impedes *TP53* expression to inhibit DNA damage-induced apoptosis"

Supplementary Table 1. Oligonucleotides used in this study

| Oligonucleotide | Sequence (5' to 3') |
|---|---|
| *ott tsa47* 545F | GACGAGATATGGGTAACGGC |
| *ott tsa47* 710R | TTCAACTGCTTCAAGTACAG |
| *TP53* 945F | TTTGGGTCTTTGAACCCTTG |
| *TP53* 1042R | CCACAACAAAACACCAGTGC |
| *MDM2* 172F | CCTGAAGATAAAGGGAAAGATA |
| *MDM2* 383R | TGGCTGCTATAAATAATGCTAC |
| *TP53* isoform 1 178F | CCCCTGTCATCTTCTGTCCC |
| *TP53* isoform 1 381R | ACATCTTGTTGAGGGCAGGG |
| *TP53* isoform 2 114F | TAGACGCCAACTCTCTCTAG |
| *TP53* isoform 2 204R | AGTCAGGGCACAAGTGAACA |
| *CDK1NA* 337F | CTGTCACTGTCTTGTACCC |
| *CDK1NA* 437R | AGTGGTAGAAATCTGTCATGC |
| *PMAIP1* 355F | CATGAGGGGACTCCTTCAAA |
| *PMAIP1* 464R | TTCCATCTTCCGTTTCCAAG |
| *SESN1* 889F | ATTCGGCTGTGGAATCAGTC |
| *SESN1* 983R | TCCACACTGTGATTGCCATT |
| *TP53I3* 114F | TAGCCGTGCACTTTGACAAG |
| *TP53I3* 241R | ACTGGCCTTGTCTCTGCATT |
| *ZMAT3* 365F | GGAACTGAAGGAGGCATCAC |
| *ZMAT3* 485R | GAATGAGCAATGTGGTCGAG |
| Used to generate and clone the coding sequences carrying the indicated amino acid substitutions into p3XFlag-CMV-7.1 | |
| *ank13* I451A F | GAACGTGGCCCCCCAGGAACTGAAAGAGAACA |
| *ank13* I451A R | TGGGGGGCCACGTTCCAGTAATTGCGTGGA |
| *ank13* I-A P452A F | CGTGGCCGCCCAGGAACTGAAAGAGAACATAC |
| *ank13* I-A P452A R | TCCTGGGCGGCCACGTTCCAGTAATTGCG |
| *ank13* IP-AA E454A F | CGCCCAGGCCCTGAAAGAGAACATACTTAAATATC |
| *ank13* IP-AA E454A R | TTCAGGGCCTGGGCGGCCACGTTCCAGTAAT |
| *ank13* IPE-AAA I459A F | AGAGAACGCCCTTAAATATCTCAATAACACGGATC |
| *ank13* IPE-AAA I459A R | TTAAGGGCGTTCTCTTTCAGGGCCTG |
| *ank13* IPEI-AAAA D467A F | TAACACGGCCCTGAGTAACATTCAGCAAGACA |
| *ank13* IPEI-AAAA D467A R | CTCAGGGCCGTGTTATTGAGATATTTAAGGGCG |

Supplementary Figure 1.

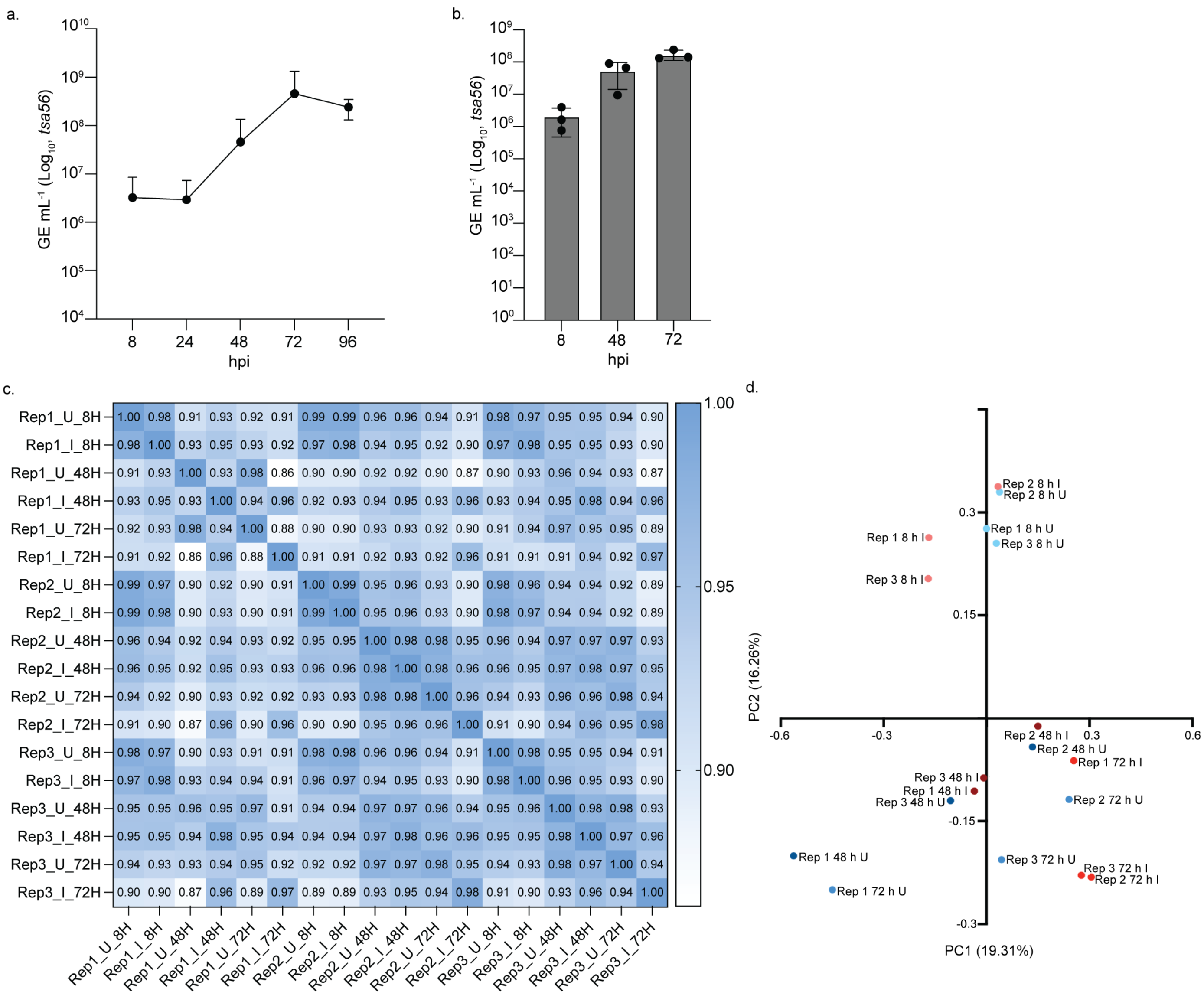


**Supplementary Fig. 1. Optimization and correlation of bulk RNAseq samples**

**(a-b)** EA.hy926 cells were infected with *O. tsutsugamushi*. GE mL$^{-1}$ was determined by qPCR for **(a)** total DNA collected from *O. tsutsugamushi* infected cells at 8, 24, 48, 72, and 96 h p.i. or **(b)** from aliquots of samples processed for RNAseq analysis and to validate consistency across replicates. Data are means $\pm$ SEM (*n* = 3 independent experiments) **(c)** A Pearson's correlation coefficient was determined for each bulk RNAseq sample individually compared to each other. An increase in blue along the gradient indicates a higher correlation. **(d)** A principal component analysis was performed. The top two principal components were plotted (PC1 and PC2). 8 h uninfected (U) samples are light blue; 8 h infected (I) are orange; 48 h U samples are navy; 48 h I samples are maroon; 72 h U samples are blue; 72 h I samples are red.

Supplementary Figure 2.

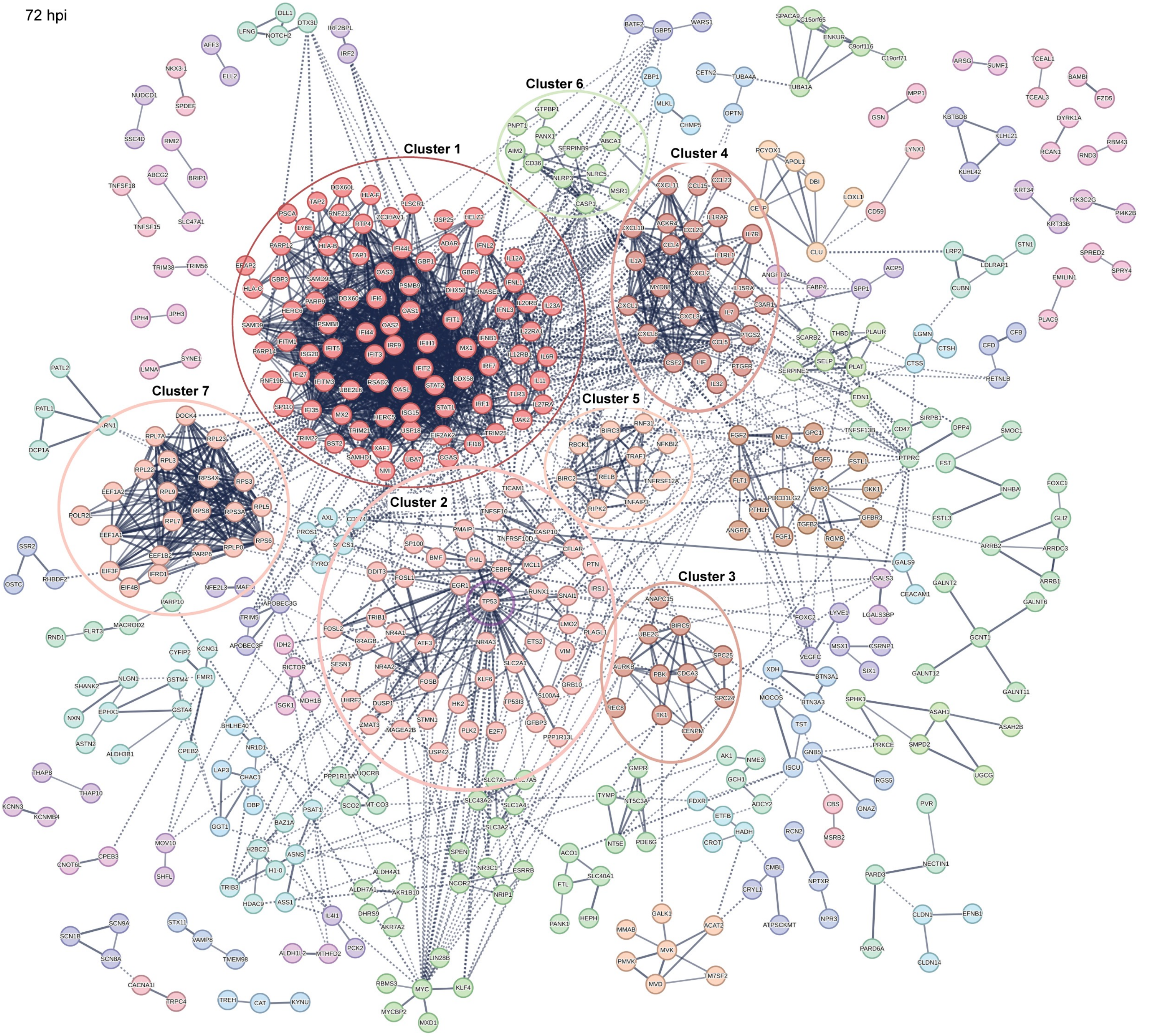


Upregulated

1 Interferon signaling (87)
2 DNA damage response, signal transduction by p53 class mediator (47)
3 Interleukin-10 signaling (25)
4 Mitotic spindle checkpoint (11)
5 TNFR1-induced NFKB signaling pathway (10)
6 Inflammasome (6)

Downregulated

7 Eukaryotic Translation Elongation (22)

**Supplementary Fig. 2. STRING analysis of DEGs at 72 h p.i.**
STRING analysis of all DEGs at 72 h p.i. Disconnected nodes were removed, and Markov Clustering was employed to define prominent clusters. Solid lines denote a direct connection between genes. Line thickness indicates the strength of the connection based on existing literature. Dotted lines represent connections between genes that are on the cusp of multiple clusters. Clusters are defined by color and cluster number. *TP53* is indicated by a purple circle.

Supplementary Figure 3.

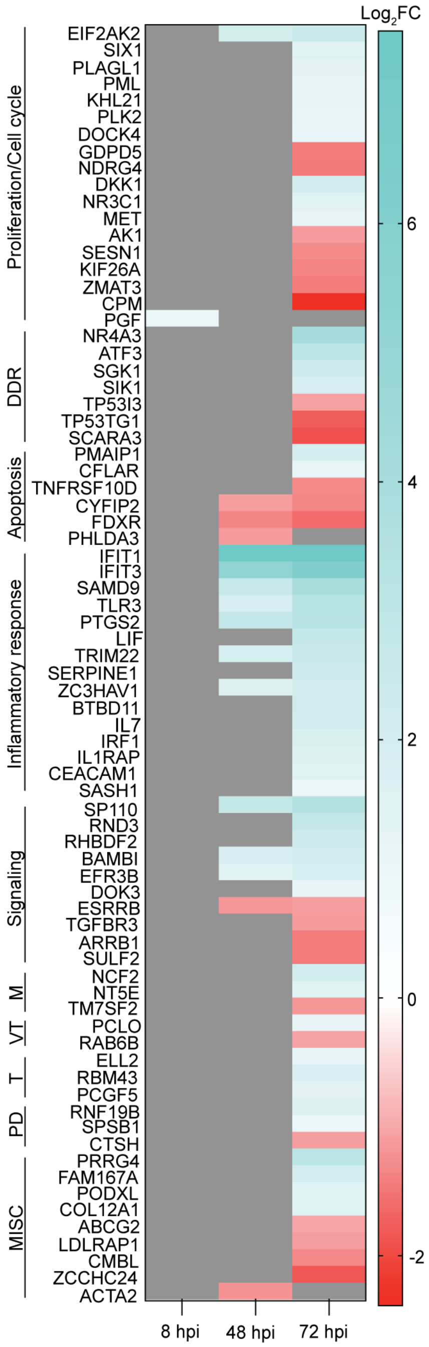


**Supplementary Fig. 3. p53 target genes are differentially expressed in *Orientia* infected cells**

A heatmap of known p53-regulated genes that are up- (teal) or downregulated (red) DEGs as defined by $\log_2$(fold change (FC)) at 8, 48, and 72 h p.i. was generated. Grey indicates no differential expression. DEGs were grouped by their roles in proliferation/cell cycle, DNA damage repair response (DDR), apoptosis, inflammatory response, signaling, metabolism (M), vesicular trafficking (VT), translation (T), protein degradation (PD), or miscellaneous processes (MISC).

Supplementary Figure 4.

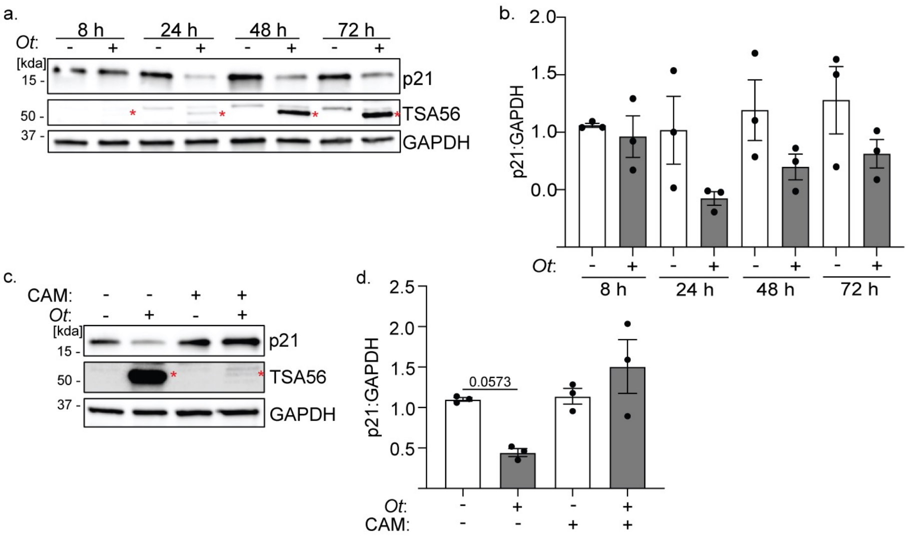


**Supplementary Fig. 4. The p53 target, p21, is downregulated in *Orientia* infected cells**
**(a-b)** WCLs of uninfected and *O. tsutsugamushi* infected samples were collected at 8, 24, 48, and 72 h p.i. and subjected to western blot analysis. **(c-d)** At 2 h p.i., uninfected and infected cells were treated with 34 ug/mL of CAM. At 72 h p.i., WCLs were analyzed by western blot. Blots were probed with p21, TSA56, and GAPDH antibodies. Red asterisks (*) denote TSA56. **(b, d)** Mean normalized ratios $\pm$ SEM of p21:GAPDH densitometric signals were calculated ($n$ = 3 independent experiments). Statistical significance was evaluated by One-way ANOVA followed by Tukey's *post hoc* test. Statistically significant values ($P \leq 0.05$) are displayed.

Supplementary Figure 5.

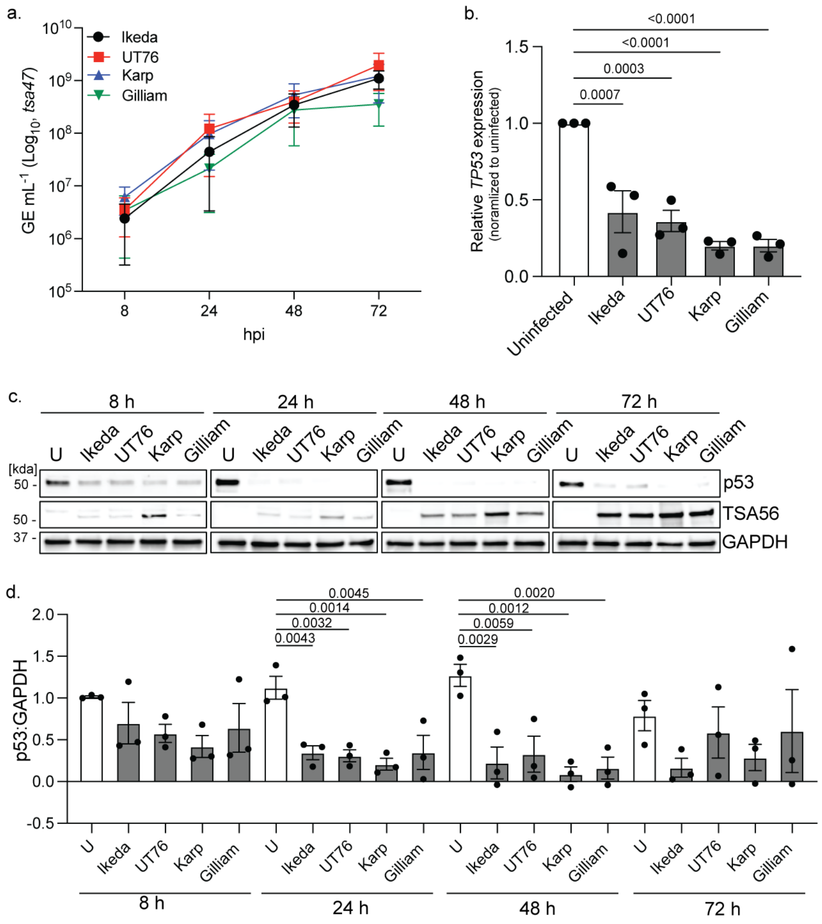


**Supplementary Fig. 5. *O. tsutsugamushi* strains reduce *TP53* and p53 expression**

**(a)** Lysates from EA.hy926 cells infected with *O. tsutsugamushi* Ikeda (black circle), UT76 (red square), Karp (blue triangle), or Gilliam (green upside-down triangle) were collected at 8, 24, 48, and 72 h p.i. GE $mL^{-1}$ was determined by qPCR of the *tsa47* gene. Data are means ± SEM ($n$ = 3 independent experiments). **(b)** RNA was collected from uninfected and Ikeda, UT76, Karp, or Gilliam infected EA.hy926 cells at 8 h p.i. Mean normalized relative expression ± SEM of *TP53* normalized to *GAPDH* was measured using RT-qPCR ($n$ = 3 independent experiments). **(c-d)** WCLs of uninfected (U) and *O. tsutsugamushi* strain infected samples collected at 8, 24, 48, and 72 h p.i. were subjected to **(c)** western blot analysis. **(d)** Mean normalized ratios ± SEM p53:GAPDH were calculated by densitometric analysis ($n$ = 3 independent experiments). Statistical significance was evaluated by One-way ANOVA followed by Tukey's *post hoc* test. Statistically significant values ($P \leq 0.05$) are displayed.

Supplementary Figure 6.

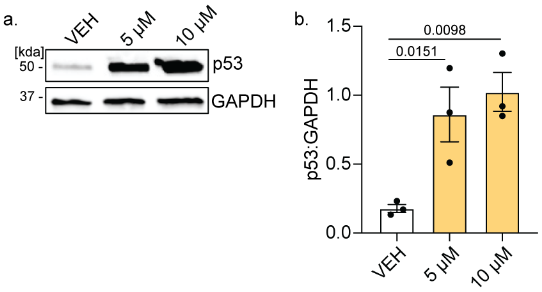


**Supplementary Fig. 6. Nutlin 3a stabilizes p53 in EA.hy926 cells**

**(a-b)** EA.hy926 cells were treated with Nutlin 3a (5 and 10 µM) for 24 h before **(a)** western blot analysis. Blots were probed with antibodies against p53, TSA56, or GAPDH. Mean normalized ratios $\pm$ SEM of **(b)** p53:GAPDH were calculated by densitometric analysis (n = 3 independent experiments). Statistical significance was evaluated by One-way ANOVA followed by Tukey's *post hoc* test. Statistically significant values ($P \leq 0.05$) are displayed.

Supplementary Figure 7.

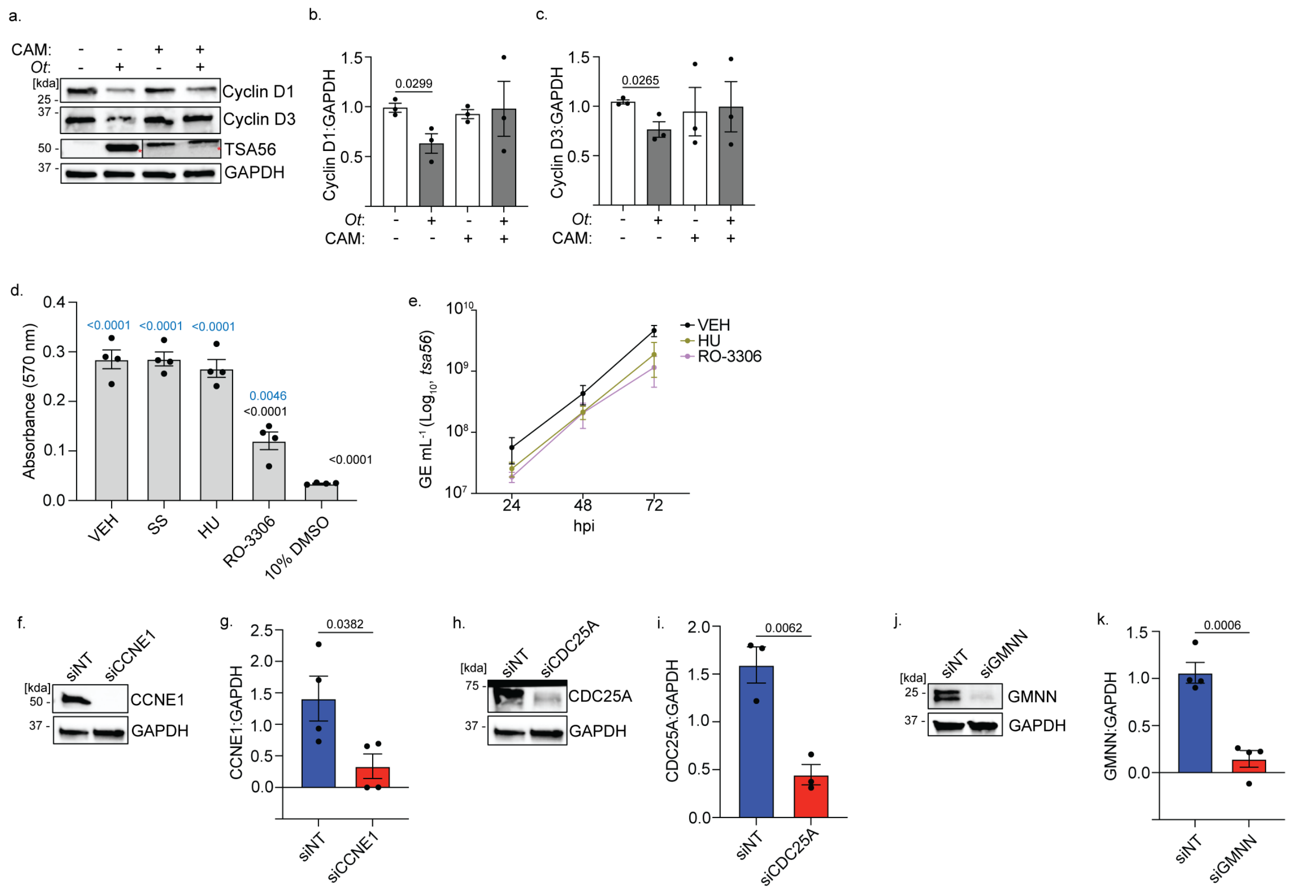


**Supplementary Fig. 7. *O. tsutsugamushi* modulates the cell cycle**

**(a-c)** At 8 h p.i., uninfected and *O. tsutsugamushi* infected samples were treated with 34 ug/mL of CAM. At 72 h p.i., WCLs were collected, and **(a)** western blot performed. Blots were probed with antibodies against **(b)** Cyclin D1 or **(c)** Cyclin D3, and TSA56 and GAPDH. Red asterisks (*) denote the TSA56. Mean normalized ratios ± SEM of Cyclin:GAPDH were calculated by densitometric analysis ($n$ = 3 independent experiments). **(d)** An MTT assay was performed on EA.hy926 cells that had been serum starved (SS), or treated with hydroxyurea (HU) or RO-3306 for 72 h. **(e)** Host cell-free *O. tsutsugamushi* was treated with HU and RO-3306 30 min prior to EA.hy926 cell inoculation. Lysates were collected at 24, 48, and 72 h p.i. and GE $mL^{-1}$ was determined by qPCR of the *tsa56* gene. **(f-k)** EA.hy926 cells were transfected with siNT, siCCNE1, siCDC25a, or siGMNN, collected at 48 h post transfection, and subjected to western blot analysis. Blots were probed with antibodies against **(f)** CCNE1, **(h)** CDC25a, or **(j)** GMNN, and GAPDH. Mean normalized densitometric ratios ± SEM of **(g)** CCNE1:GAPDH, **(i)** CDC25a:GAPDH, or **(k)** GMNN:GAPDH were calculated ($n$ = 3-4 independent experiments). Statistical significance was evaluated by One-way ANOVA followed by Tukey's *post hoc* test or an unpaired student's *t*-test. Statistically significant values ($P \leq 0.05$) are displayed.

Supplementary Figure 8.

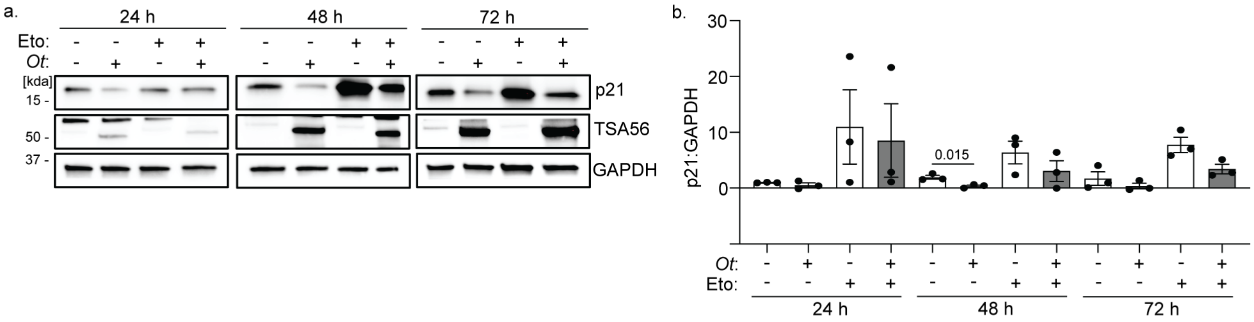


**Supplementary Fig. 8. Etoposide does not rescue p21 levels in *Orientia* infected cells**

**(a, b)** Uninfected and infected EA.hy926 were treated with etoposide (Eto) 24 h prior to collection at 24, 48, and 72 h p.i. **(a)** WCLs were collected and subjected to western blot analysis. Blots were probed for p21, TSA56, and GAPDH. Red asterisks (*) denote TSA56 in infected samples. Mean normalized ratios $\pm$ SEM of **(b)** p21:GAPDH were calculated by densitometric analysis ($n$ = 3 independent experiments). Statistical significance was evaluated by One-way ANOVA followed by Tukey's *post hoc* test. Statistically significant values ($P \leq 0.05$) are displayed.

Supplementary Figure 9.

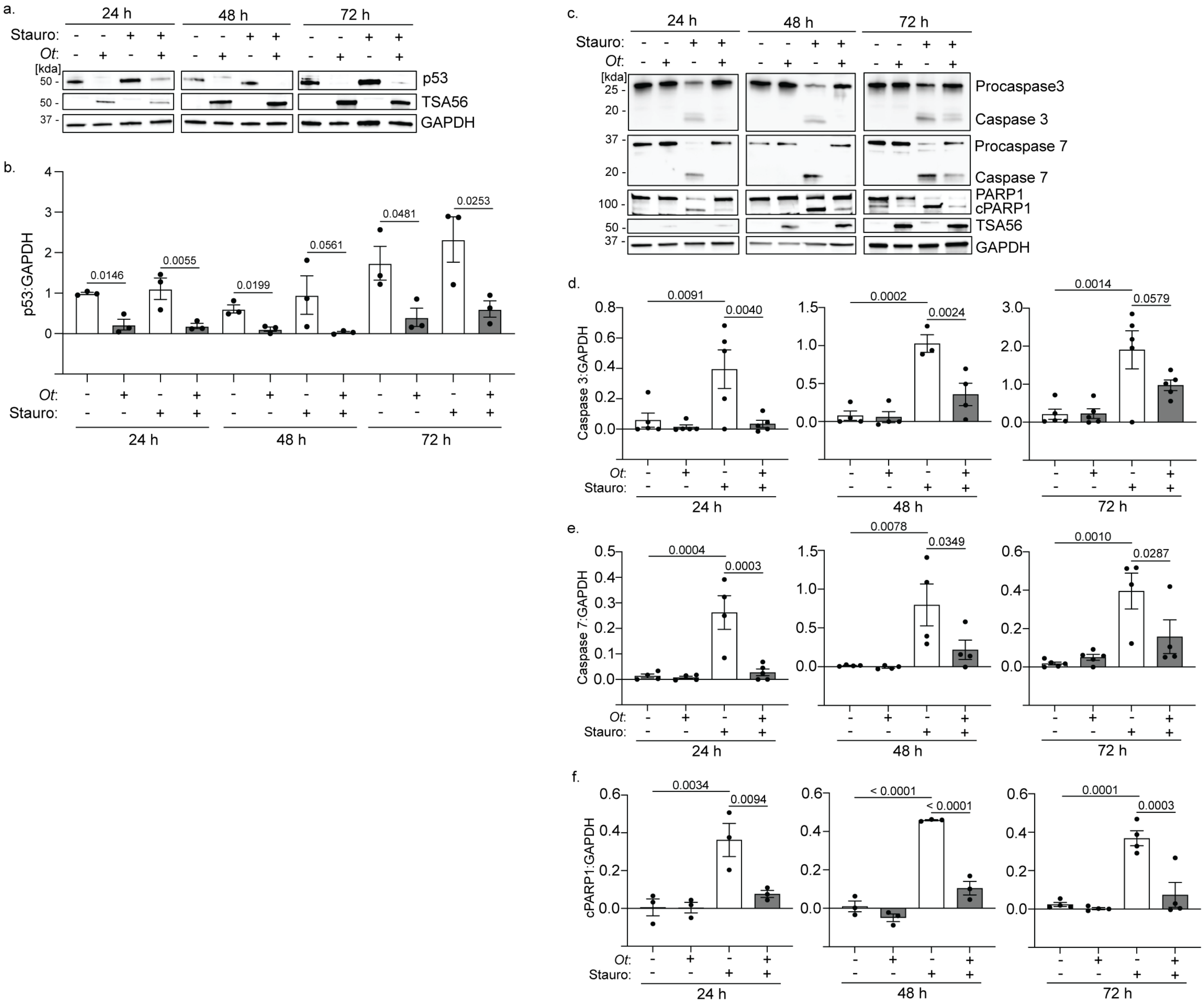


**Supplementary Fig. 9. *O. tsutsugamushi* prevents staurosporine-induced apoptosis**

**(a-d)** Uninfected and infected EA.hy926 were treated with staurosporine (stauro; 1 μM) 24 h prior to collection. **(a, c)** WCLs were collected at 24, 48, and 72 h p.i. and subjected to western blot analysis. Blots were probed for p53, procaspase 3 and cleaved caspase 3, procaspase 7 and cleaved caspase 7, full length and cleaved (cPARP1) PARP1, TSA56, and GAPDH. Red asterisks (*) denote TSA56 in infected samples. Mean normalized densitometric ratios ± SEM of each **(b)** p53:GAPDH, **(d)** Caspase3:GAPDH, **(e)** Caspase7:GAPDH, and **(f)** cPARP1:GAPDH were calculated ($n$ = 3 independent experiments). Statistical significance was evaluated by One-way ANOVA followed by Tukey's *post hoc* test. Statistically significant values ($P \leq 0.05$) are displayed.

Supplementary Figure 10.

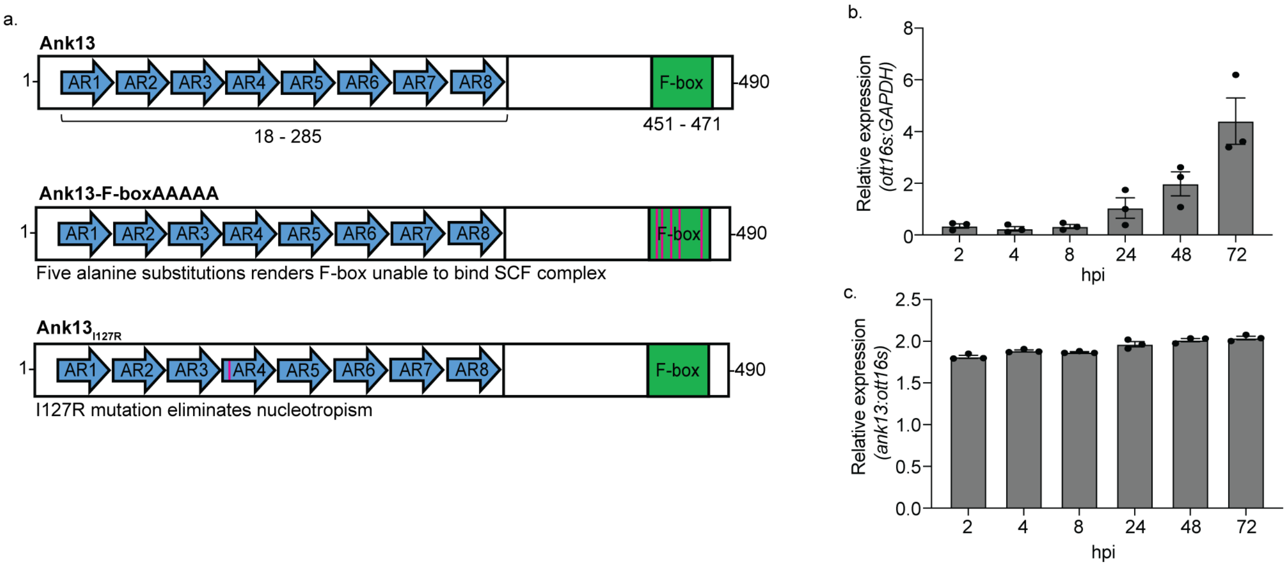


**Supplementary Fig. 10. *O. tsutsugamushi* expresses *ank13* in endothelial cells**
**(a)** Schematic depicting Ank13 and its mutants, Ank13-F-boxAAAAA and $\text{Ank13}_{\text{I127R}}$. The eight ankyrin repeats are displayed as blue arrows. The F-box is green. The five alanine substitutions in Ank13-F-boxAAAAA and the I127R mutation in $\text{Ank13}_{\text{I127R}}$ are presented as red bars. **(b-c)** RNA was collected from *O. tsutsugamushi* infected EA.hy926 cells at 2, 4, 8, 24, 48, and 72 h p.i.. RT-qPCR was performed to determine relative gene expression of **(b)** *ott16s* normalized to *GAPDH* and **(c)** *ank13* normalized to *ott16s* (*n* = 3 independent experiments).
